# Distinct macrophage and T cell programs shape pancreatic inflammation during metabolic stress and aging

**DOI:** 10.64898/2026.07.08.737054

**Authors:** Somesh Sai, Ibrahim Omar, Matthias Barone, Kerstin Mühle, Maria Schneider, Fenfen Liu, Sharanya Sriram, Juliette Claire Johnson, Tizia Thoma, Thomas Conrad, Tatiana Borodina, Birgit Sawitzki, Maike Sander, Han Zhu

## Abstract

Type 2 diabetes is linked to systemic inflammation driven by metabolic stress and aging. Although pancreatic inflammation associated with these factors is well documented, the dynamics of immune cell populations and their molecular changes remain poorly understood. We characterized immune cell alterations in the pancreas and pancreatic islets during Western diet (WD) feeding and aging using imaging mass cytometry (IMC) and single-cell RNA sequencing (scRNA-seq). Spatial and transcriptional analyses were performed to define immune cell subtype composition, activation states, and inferred cell-cell communication programs under metabolic and age-related stress conditions. Our analyses identified expansion of an F4/80^low^ macrophage subtype and activated effector-like CD8^+^ T cells throughout the pancreas during WD feeding and aging. Within pancreatic islets, single-cell RNA sequencing identified a type I interferon-responsive macrophage population with low F4/80 expression that expanded during overnutrition. Notably, the type I interferon responses elicited by these stressors diverged: aging was associated with a more canonical type I interferon response, whereas overnutrition induced a broader response that included STAT3-associated transcriptional programs. We further provide evidence for enhanced cytokine-mediated communication between macrophages and a CD8^+^ cytotoxic T-cell population under overnutrition and aging. These findings show that metabolic stress and aging remodel pancreatic inflammation through overlapping but distinct immune mechanisms, involving expansion of F4/80^low^ macrophages, activation of divergent type I interferon programs, and enhanced macrophage–CD8^+^ T-cell communication. Together, these findings suggest that distinct therapeutic approaches may be required to preserve islet function in type 2 diabetes driven by metabolic stress versus aging.

**Article Highlights:**

- Metabolic stress and aging remodel pancreatic inflammation through overlapping but distinct immune mechanisms.
- Spatial and single-cell analyses identified conserved remodeling of pancreatic macrophage populations accompanied by activation of cytotoxic T-cell responses during metabolic stress and aging.
- In pancreatic islets, macrophages exhibited distinct type I interferon-associated transcriptional programs, with aging showing a more canonical interferon response and metabolic stress eliciting a broader inflammatory program that included STAT3-associated transcriptional signatures.
- These findings provide a framework for understanding how metabolic stress and aging differentially shape pancreatic inflammation.

## INTRODUCTION

Metabolic disorders such as type 2 diabetes (T2D) are on the rise worldwide [1]. Obesity and aging are major risk factors for T2D, both commonly linked to systemic, low-grade inflammation [2–6]. Inflammation in adipose tissue, liver, and skeletal muscle promotes insulin resistance and hyperglycemia [7, 8]. In T2D, inflammation also affects the pancreatic islets, where insulin-producing beta cells reside [9]. This islet inflammation is characterized by increased immune cell infiltration and elevated pro-inflammatory cytokines, which impair beta cell function and reduce insulin secretion - critical features in T2D pathogenesis [10, 11]. It remains unclear whether islet inflammation is predominantly triggered by obesity or aging, and whether the resulting inflammatory patterns differ between these risk factors.

Islet inflammation in obesity is associated with an increase in myeloid-lineage cells, including monocytes and macrophages [12, 13]. Increased macrophage numbers in islets have been observed in both high-fat diet-induced models [10] and genetically modified (*db/db*) overnutrition models [14]. This inflammation has been associated with beta cell dysfunction, with current research focusing on uncovering the origins and phenotypic characteristics of the macrophages involved. Obesity-related islet inflammation has been shown to involve contributions from both monocyte-derived macrophages [15] and islet-resident macrophages [16]. These macrophages are characterized by enhanced chemokine production and exhibit a pro-inflammatory phenotype resembling M1 macrophage-like properties [14]. However, transcriptomic analyses suggest that these immune cells do not align precisely with the traditional M1/M2 macrophage polarization paradigm [17]. This likely reflects the heterogeneity of macrophage subpopulations within islets, with distinct subsets exhibiting different responses to metabolic stressors such as overnutrition. T cells also contribute to obesity-associated inflammation in metabolically active tissues by promoting cytokine production and macrophage activation [18, 19]. However, the involvement of adaptive immune cells in obesity-related islet inflammation remains poorly understood. This knowledge gap is partly due to the low abundance of these immune cells in islets [13].

Aging, a major risk factor for T2D, is also associated with low-grade chronic inflammation, commonly referred to as inflammaging. Aging-associated islet inflammation has been documented across multiple species, including zebrafish [20], mice [21], rats [22], and humans [23]. This inflammatory state impairs beta cell proliferative capacity, primarily through elevated production of TNFα [20], and contributes to beta cell dysfunction via activation of the Toll-like receptor-4 pathway [21]. Notably, there is a significant increase in T cell infiltration in aging islets [23]. These findings highlight distinct inflammatory responses between obesity-related and aging-related islet dysfunction, underscoring the need for further investigation into their mechanisms and interactions.

Advances in single-cell transcriptomics and high-throughput spatial-omics have transformed the profiling of complex inflammatory mechanisms at cellular and spatial levels [24, 25]. These technologies have been applied to investigate cellular changes in the pancreas and islets in type 1 diabetes (T1D) [26–29] and T2D [30, 31]. However, the low abundance of immune cells within islets has hindered comprehensive analyses of immune cell dynamics in metabolic diseases, particularly in T2D, where islet inflammation evolves gradually rather than undergoing rapid changes over short periods.

In this study, we combined imaging mass cytometry (IMC) and single-cell RNA sequencing (scRNA-seq) to define immune cell composition, transcriptional states, and spatial organization in the pancreas and pancreatic islets during Western diet (WD) feeding and aging. Our analyses reveal that metabolic stress and aging induce overlapping yet distinct inflammatory programs. Both conditions remodel pancreatic macrophage and T-cell populations, whereas islet macrophages exhibit context-dependent type I interferon–associated transcriptional programs, with aging characterized by a more canonical interferon response and WD feeding eliciting a broader inflammatory program that includes STAT3-associated transcriptional signatures. Together, these findings provide a framework for understanding how metabolic stress and aging differentially shape pancreatic inflammation and may contribute to beta cell dysfunction.

## RESEARCH DESIGN AND METHODS

### Data and Resource Availability

Single-cell RNA sequencing data have been deposited in the Gene Expression Omnibus (GEO) under accession number GSE266277 and will be made publicly available upon publication. Processed datasets used for analysis will be deposited in Zenodo, and analysis code used to generate figures will be deposited on GitHub prior to publication. Additional information required to reanalyze the data is available from the corresponding author upon reasonable request. No unique reagents, animal models, antibodies, viral vectors, or other non-commercial resources were generated in this study.

### Mice husbandry

Male C57BL/6J mice (10 and 78 weeks old; The Jackson Laboratory, no. 000664) were housed in an Association for Assessment and Accreditation of Laboratory Animal Care–accredited facility at the University of California, San Diego under protocols approved by the Institutional Animal Care and Use Committee. Mice were maintained on a 12-hour light/dark cycle with ad libitum access to water and standard rodent chow (PicoLab Rodent Diet 20 5053). To establish the aging cohort, 78-week-old mice were maintained under these conditions for an additional 27 weeks. To induce metabolic stress, mice were fed a Western diet (TD.88137; Envigo Teklad) containing 42% of calories from fat and 42.7% from carbohydrates.

### Imaging mass cytometry

For IMC analysis, pancreatic tissues from a minimum of four mice per experimental condition and time point were processed. At least 10 regions of interest (ROIs) were analyzed for each condition, resulting in a total of 78 ROIs. ROIs were randomly distributed across 17 tissue microarrays (TMAs), with each TMA containing samples from multiple experimental groups and time points to minimize batch effects. Antibodies were conjugated to metal isotopes using the Maxpar X8 antibody labeling kit (Fluidigm), titrated prior to use, and are listed in Supplementary Table 1B. TMA sections were stained with a panel of metal-conjugated antibodies using a standard IMC staining protocol with Iridium DNA intercalator, followed by image acquisition on a Hyperion Tissue Imager coupled to a Helios mass cytometer (Fluidigm).

### Fluorescence-activated cell sorting (FACS) of CD45^+^/CD45^−^ islet cells

Pancreatic islets were isolated using a modified collagenase digestion protocol. Briefly, Collagenase P (Roche, #11249002001; 0.8 mg/mL) was perfused through the common hepatic bile duct, and pancreata were digested at 37°C for approximately 15 min. Islets were purified by HBSS/Histopaque density-gradient separation followed by two rounds of manual handpicking under a dissection microscope to minimize exocrine contamination. Immediately after isolation, islets were dissociated into single-cell suspensions using Accumax (Invitrogen, 00-4666-56) for 12 min at 37°C. Cells were washed, stained with Alexa Fluor 488-conjugated anti-CD45 antibody (1:300) for 30 min at 4°C, followed by Aqua Dead Cell Stain (Invitrogen, L34957), and sorted into viable CD45^+^ and CD45^−^ populations using a FACSAria II Cell Sorter (BD Biosciences). Islets from at least 20 mice were pooled for each sample to obtain sufficient CD45+ cells for single-cell RNA sequencing.

### Single-cell RNA-sequencing (scRNA-seq)

Sorted cells were pelleted (250 × g, 5 min, 4°C), counted using a Scepter automated cell counter, and 10,000 cells per sample were loaded onto a 10x Chromium Controller for GEM generation and cell barcoding using Next GEM Single Cell 3′ v3.1 reagents. Barcoded cells were processed for cDNA synthesis and library construction according to the manufacturer’s instructions. Libraries were quantified using a Qubit fluorimeter (Life Technologies), fragment size was verified using a TapeStation (High Sensitivity D1000; Agilent), and sequencing was performed on Illumina NextSeq 500, HiSeq 4000, or NovaSeq 6000 instruments. Sequencing reads were trimmed prior to downstream analysis.

### Metabolic studies

For glucose tolerance tests (GTT), mice were fasted for 6 h before intraperitoneal (i.p.) injection of glucose (1.5 mg/g body weight). Blood glucose was measured from tail vein blood at baseline and 20, 40, 60, 90, 120, and 150 min using a Bayer Contour glucometer. For insulin tolerance tests (ITT), mice were fasted for 5 h before i.p. injection of insulin (0.8–2.0 U/kg body weight), and glucose was monitored as described for GTT. Plasma insulin concentrations were measured in 6-h-fasted mice before and 10 min after i.p. glucose injection (1.5 mg/g body weight) using Mouse Ultrasensitive Insulin ELISA or Mouse Insulin ELISA kits (ALPCO).

### Glucose-stimulated insulin secretion

Glucose-stimulated insulin secretion (GSIS) assays were performed as previously described [35]. Islets isolated from six to ten mice per group were pooled and cultured overnight in RPMI 1640 containing 8 mM glucose and standard supplements. Islets were then pre-incubated for 1 h in Krebs-Ringer bicarbonate HEPES (KRBH) buffer containing 2.8 mM glucose before groups of 10 size-matched islets were transferred to KRBH containing either 2.8 or 16.8 mM glucose for 1 h at 37°C with 5% CO₂. Supernatants were collected, and islets were lysed overnight in 20% acid/80% ethanol. Insulin concentrations in supernatants and lysates were measured using a mouse insulin ELISA (ALPCO). Insulin secretion was expressed as the percentage of total insulin content per hour to account for variation in islet size.

### Imaging mass cytometry data analysis

IMC images were processed using a standardized single-cell analysis workflow that included image preprocessing, cell segmentation, marker normalization, batch correction, and immune cell identification. Single-cell marker intensities were used for unsupervised Phenograph clustering and UMAP visualization to define immune cell populations. Pancreatic islets were identified based on insulin, NKX6-1, and GLP1R staining, allowing classification of cells into exocrine, peri-islet, and islet regions. Immune cell densities were quantified within each compartment and across the whole pancreas.

### Single-cell RNA-sequencing data analysis

Raw sequencing data were processed using Cell Ranger followed by ambient RNA correction, quality-control filtering, doublet removal, normalization, batch integration, dimensionality reduction, and unsupervised clustering. Immune and non-immune cell populations were annotated using canonical marker genes, and macrophage, T-cell, and beta-cell populations were subsequently subsetted for downstream analyses. Differential gene expression analysis was performed using the LIBRA package with Seurat-based statistical testing. Differentially expressed genes were grouped by K-means clustering to identify gene modules with shared expression dynamics, and functional enrichment was assessed using Enrichr. Cell-cell communication between macrophage and T-cell populations was inferred using CellChat based on curated ligand–receptor interactions, and differential interaction analyses were performed between experimental conditions.

### Quantification and statistical analysis

Statistical analyses were performed using GraphPad Prism (v8.1.2) and R (v3.6.1). Sample sizes (n), statistical tests, and P values are provided in the figures and figure legends. For metabolic studies, n represents biological replicates (individual mice). Data are presented as mean ± SD. One-way or two-way ANOVA with appropriate multiple-comparison tests was used as indicated.

## RESULTS

### Spatial profiling of pancreatic immune cell dynamics during Western diet feeding and aging

To examine the effects of WD feeding and aging on pancreatic immune cells, we used IMC to analyze tissue sections from mice on WD or chow diets for 1, 12, and 24 weeks, as well as from aged mice (105-week-old) to compare metabolic- and aging-related changes (**Figure 1A and Supplementary Table 1A**). In this analysis, the effects of metabolic stress were assessed by comparing WD-fed mice with time-matched chow-fed controls, while the effects of aging were evaluated by comparing the aged group to a reference group (“adult”) aggregated from all chow-fed cohorts across various feeding times examined.

**Figure 1:**
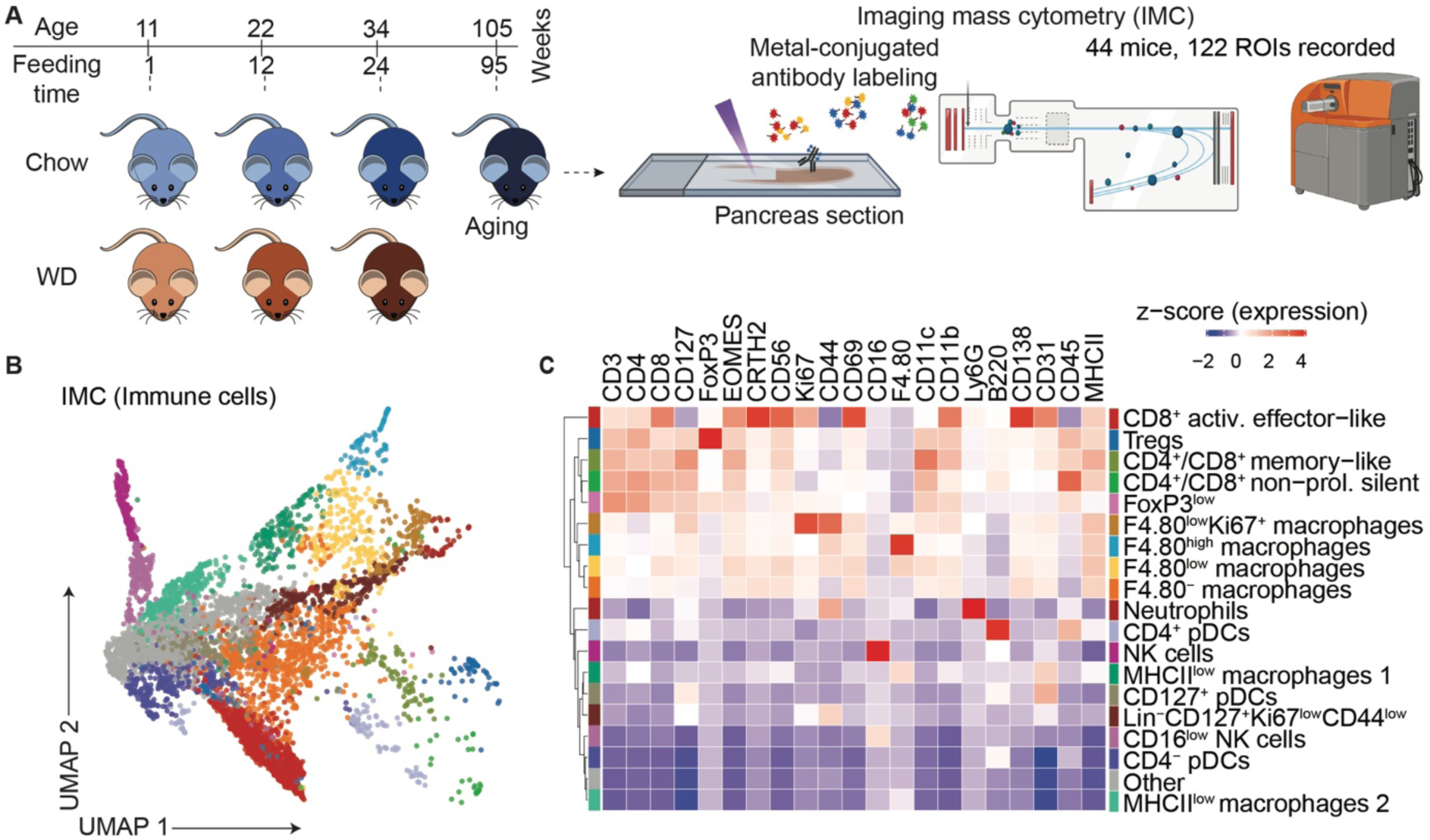
Imaging mass cytometry (IMC) characterization of pancreatic immune cell dynamics during Western diet (WD) feeding and aging. **(A)** Spatial analysis experimental design. Pancreas samples were harvested from four mice at 1, 12, and 24 weeks of WD or chow diet feeding. Aged samples were obtained from 2-year-old mice. Samples were analyzed by IMC to profile immune and islet-specific marker proteins. **(B)** UMAP embedding of CD45^+^CD19^-^ immune cells identified by IMC. Cell identities were annotated based on marker protein expression shown in (**C**). **(C)** Heatmap of z-scored immune cell marker protein expression across CD45^+^CD19^-^ immune cell subtypes.

IMC analysis was performed on tissue microarrays containing samples from all experimental conditions using a panel of 25 immune and islet markers (**Supplementary Table 1B**). Following cell segmentation and quality control (**Supplementary Figure S1A–C**), CD8^+^ and CD11c^+^ cells were readily detected across all samples (**Supplementary Figure S1D**). Because CD19^+^ B cells primarily localized to pancreatic lymph nodes (**Supplementary Figure S1E**), they were excluded from downstream analyses. Phenograph [32] clustering of CD45^+^CD19^−^ immune cells identified multiple immune populations (**Figure 1B,C**), including activated effector-like CD8+ T cells (Eomes^+^CD56^+^Ki67^+^CD69^+^), regulatory T cells (Foxp3^+^), memory-like T cells (CD127^high^CD4^+^CD8^+^), non-proliferative T cells (Ki67^−^CD4^+^CD8^+^), Foxp3^low^CD4^+^ T cells, six macrophage populations distinguished by F4/80, CD11c, Ki67, and MHCII expression, three B220^+^ plasmacytoid dendritic-like cell populations (CD4^+^, CD4^−^, and CD127^+^), Ly6G^+^ neutrophils, CD16^high^ NK cells, and Lin^−^CD127^+^ innate lymphoid-like cells.

### Metabolic stress induces accumulation of inflammatory F4/80^low^ macrophages and CD8^+^ activated effector-like T-cells in the pancreas

We next examined changes in pancreatic immune cell abundance under WD feeding and aging by quantifying immune cell proportions and absolute cell densities using IMC (**Supplementary Table 1C**). Under metabolic stress, we observed increased proportions of F4/80^high^, F4/80^low^, and F4/80^-^ macrophages as well as CD8^+^ activated effector-like T-cells under metabolic stress. Regions of interest were further classified as exocrine, peri-islet, or islet based on insulin, NKX6-1, and GLP1R staining (**Figure 2A**).

**Figure 2:**
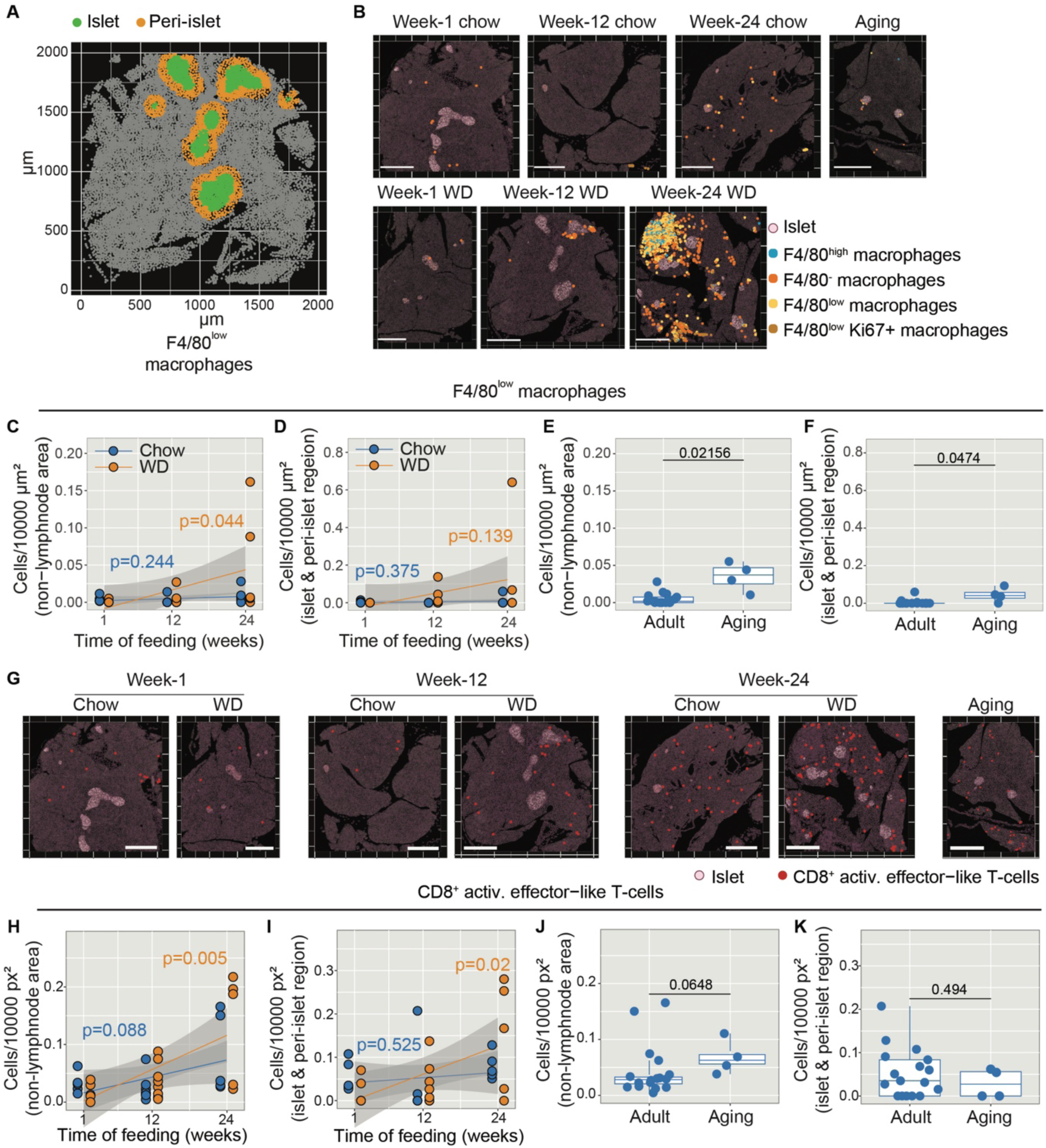
Metabolic stress enhances age-driven accumulation of inflammatory F4/80low macrophages and CD8^+^ activated effector-like T-cells in the pancreas. **(A)** Representative image showing pancreatic islets (green) and their peri-islet regions (yellow) defined as the area extending 70 μm beyond the islet boundary. Axes are labeled in μm to indicate spatial dimensions. **(B)** Representative ROIs showing the three macrophage subtypes (F4/80^low^ Ki67^+^, F4/80^high^, F4/80^low^, and F4/80^-^) characterized by different F4/80 levels. Pancreatic islet regions are indicated by raw insulin signal (white). Scale bar, 500 μm. (**C,D**) Scatter plots depicting the cell counts of F4/80^low^ macrophages per 10000 μm^2^of the pancreas (excluding lymph nodes, **C**) and per 10000 μm^2^ of the islets and peri-islet regions (**D**) across different time points and feeding conditions. Solid lines represent linear regression fits, with shaded areas indicating 95% confidence intervals calculated from the standard error. Each condition includes data from n ≥ 3 mice. P-values for the regression coefficient slope were determined using a linear model and t-tests. (**E,F**) Box plots showing the cell counts of F4/80^low^ macrophages per 10000 μm^2^ of the pancreas (excluding lymph nodes, **E**) and per 10000 μm^2^ of islets and peri-islet regions (**F**) in adult and aging mice. Each condition includes data from n ≥ 3 mice. P-values were determined by Wilcoxon signed-rank test with Benjamini & Hochberg adjustment. (**G**) Representative ROIs showing CD8^+^ activated effector-like T-cells in the pancreas. Pancreatic islet regions are marked by raw insulin signal (white). Scale bar 500 μm. (**H,I**) Scatter plot showing the cell counts of CD8^+^ activated effector-like T cells per 10000 μm^2^ of the pancreas (excluding lymph nodes, **H**) and per 10000 μm^2^ of the islets and peri-islet regions (**I**) across various time points and feeding conditions. Solid lines represent linear regression fits, with shaded areas indicating 95% confidence intervals calculated from the standard error. Each condition includes data from n ≥ 3 mice. P-values for the regression coefficient slope were determined using a linear model and t-tests. (**J,K**) Box plots showing the cell counts of CD8^+^ activated effector-like T cells per 10000 μm^2^ of the pancreas (excluding lymph nodes, **J**) and per 10000 μm^2^ of islets and peri-islet regions (**K**) in adult and aging mice. Each condition includes data from n ≥ 3 mice. P-values were determined by Wilcoxon signed-rank test with Benjamini & Hochberg adjustment.

We first analyzed macrophage subtypes with different levels of F4/80 expression (**Figure 2B**). Among these subtypes, only F4/80^low^ macrophages exhibited a feeding time–dependent increase across the pancreas under WD feeding, an effect not observed under chow conditions (**Figure 2C**). Notably, this expansion was restricted to the exocrine compartment and was not detectable within islets or peri-islet regions (**Figure 2D**). F4/80^low^ macrophages also accumulated in the aged pancreas (**Figure 2E**), and in contrast to the WD condition, this increase extended into regions proximal to pancreatic islets (**Figure 2F**). We did not observe significant changes in the density of other macrophage subtypes (**Supplementary Figure S2A-L**) under either WD or aging condition. These observations suggest that macrophage numbers increase in the pancreas under both metabolic stress and aging, and that this accumulation happens in a subtype-specific manner.

Similarly, we quantified the density of CD8+ activated effector-like T-cells across different conditions (**Figure 2G**). We observed a notable expansion of these T-cells in response to WD feeding, both throughout the pancreas and within islet and peri-islet regions (**Figure 2H,I**). In contrast to the F4/80^low^ macrophage subtype, the CD8+ activated effector-like T-cells did not significantly accumulate in aged pancreas (**Figure 2J,K**). Taking advantage of the spatial information provided by IMC, we next examined whether any macrophage subtypes co-localized with CD8^+^ activated effector-like T cells by assessing correlations in their absolute cell numbers within the same ROIs. We found positive spatial correlations between CD8^+^ activated effector-like T cells and F4/80^low^ macrophages, and an even stronger correlation with F4/80^-^ macrophages (**Supplementary Figure S2M**). Interestingly, F4/80^-^ macrophages did not accumulate in the pancreas under any of the conditions tested (**Supplementary Figure S2C,F,I,L**), suggesting that their spatial association with CD8^+^ activated effector-like T cells may reflect a signaling or interaction role.

### Single-cell profiling reveals the islet immune landscape

Due to the scarcity of immune cells in pancreatic islets, our IMC analysis did not allow us to investigate islet inflammation under metabolic stress and aging conditions. To complement the spatial data, we performed scRNA-seq on FACS-sorted CD45^+^ immune and CD45^-^ non-immune cells from isolated islets of the same cohorts. With the goal of capturing early molecular alterations induced by WD feeding, we focused our scRNA-seq analysis on the 1-week and 12-week time points (**Figure 3A**).

**Figure 3.**
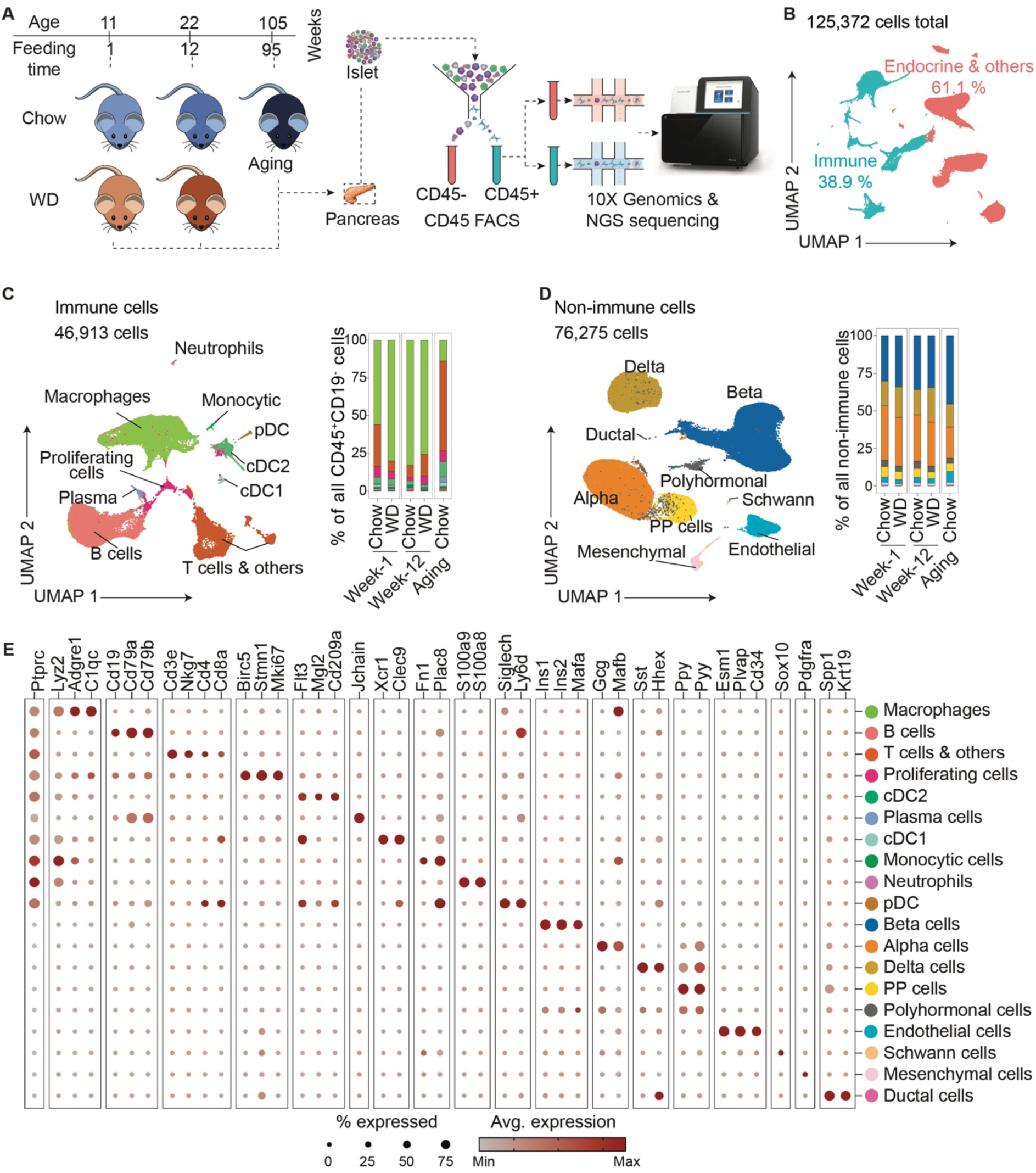
Single-cell RNA-sequencing analysis of pancreatic islet associated immune cells during WD feeding and aging. **(A)** Single-cell transcriptome experimental design. Pancreas samples were harvested from four mice at 1, and 12 weeks of WD or chow diet feeding. Aged samples were obtained from 2-year-old mice. Islets were collected from 12 cohorts among all conditions (1- and 12-week WD or chow diet, and aging), each cohort comprising over 20 mice. CD45^+^ immune cells and CD45- non-immune cells were processed separately for scRNA-seq. **(B)** UMAP embedding of 125,372 high-quality single-cell transcriptomes across 12 cohorts. Cells were classified as immune (CD45^+^; *Ptprc*) or non-immune. Proportions of annotated populations are shown. **(C)** UMAP embedding of immune cells from panel **B**, with clusters identified by known immune markers. The relative abundance of immune cell types, excluding B cells, under each condition is shown (right). **(D)** UMAP embedding of non-immune cells from panel **B**, with clusters annotated by known gene markers. The relative abundance of each cell type under each condition is shown (right). **(E)** Dot plot of hallmark genes for cell types identified in panels **C** and **D**.

After quality control (**Supplementary Figure S3A–C and Supplementary Table 1D**), 125,372 high-quality single-cell transcriptomes were retained, including 46,913 CD45^+^ immune cells and 76,275 CD45^−^ non-immune cells (**Figure 3B–D**). Marker gene analysis identified macrophages (*Lyz2, Adgre1, C1qc*), monocytes (*Fn1, Plac8*), neutrophils (*S100a9, S100a8*), dendritic cells(*Xcr1, Flt3, Siglech*), T cells (*Cd3e, Cd4, Cd8a*), B cells (*Cd19, Cd79a, Cd79b*), plasma cells (*Jchain*), and major non-immune cell populations (**Figure 3E**). Similar to the IMC analysis, B cells and polyhormonal endocrine cells were excluded from downstream analyses (**Supplementary Figure S3D,E**). Macrophages represented the predominant immune population in adult islets, whereas aged islets showed an increased proportion of T cells (**Figure 3C**), consistent with previous studies [23].

### Distinct macrophage interferon responses to western diet feeding and aging

We next focused on phenotypic changes in the macrophage populations under metabolic stress and aging. To investigate macrophage heterogeneity within islets, we performed sub-clustering of macrophages from our scRNA-seq dataset (**Figure 4A**) and identified five distinct subpopulations. Macs-1 exhibited relatively high F4/80 (Adgre1) and low CD11c (Itgax)/CD69 (Cd69) expression, consistent with a homeostatic state, whereas Macs-2–5 displayed progressively lower F4/80 and higher CD11c/CD69 expression, indicative of activated states. Macs-4 uniquely expressed high Ki67 (Mki67), consistent with a proliferative phenotype (**Figure 4B**). Macs-2, Macs-3, and Macs-5 resembled the previously described Atf3^+^, Cxcl9^+^, and Prdx1^+^ macrophage states in NOD mouse islets (**Supplementary Figure S4A**) [29].

**Figure 4:**
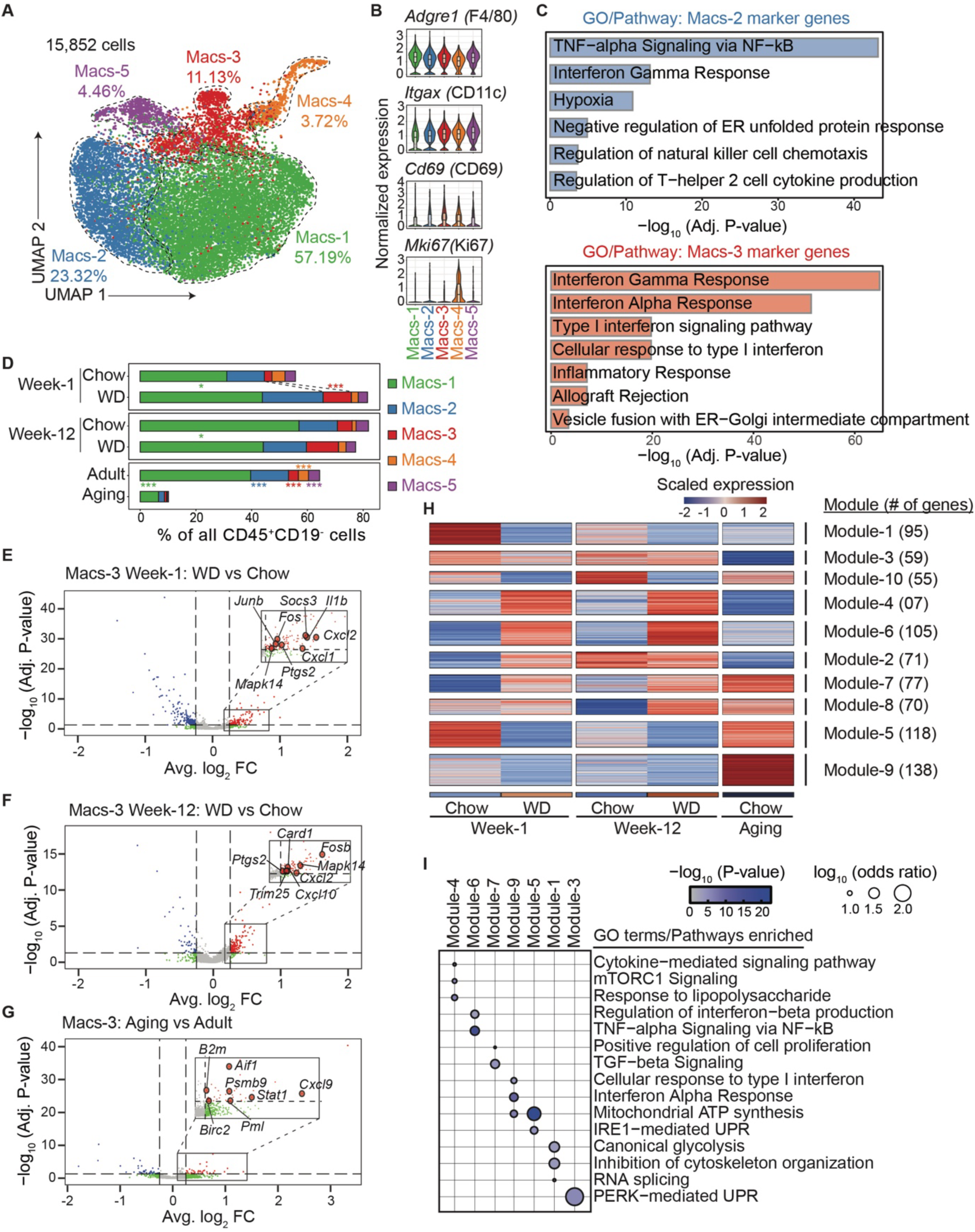
Aging and metabolic stress activate distinct type I interferon responses in inflammatory macrophages. **(A)** UMAP embedding of islet-associated macrophages pooled across time points (week-1, week-12, and aging) and experimental conditions (WD and chow). The overall proportion of each defined subtype is indicated. **(B)** Violin plots showing normalized expression levels of genes of interest across macrophage subpopulations. **(C)** Enriched gene ontology (GO) terms for marker genes in Macs-2 (top) and Macs-3 (bottom). Significance (-log_10_ adjusted p-value) of the enrichments is shown. **(D)** Proportions of macrophage subpopulations expressed as percentages of all immune cell types, excluding B cells, in each experimental group. Data were pooled across replicates for each group. * p<0.05, *** p < 0.001, calculated using a mixed-effects binomial model. **(E-G)** Volcano plots showing differentially expressed genes in Macs-3 under different conditions: (**E**) WD versus chow at week-1, (**F**) WD versus chow at week-12, and (**G**) aging versus adult (combined week-1 and week-12 chow data). Genes with significant upregulation and downregulation are highlighted in red and blue, respectively. Dashed vertical lines indicate an absolute average log_2_-fold change threshold of 0.25, while the dashed horizontal line represents a significance threshold of –log_10_(adjusted p-value) = 1.3 (corresponding to adjusted p-value of 0.05). Genes involved in type I interferon response are highlighted. **(H**) K-means clustering of differentially expressed genes in Macs-3 across all possible pairwise combinations across the 5 experiment groups. The color bar represents the scaled gene expression levels across all conditions analyzed. **(I)** Dot plot showing enriched gene ontology (GO) terms for selected k-means gene modules from **(H)**. Significance (-log_10_ p-value) and log_10_ odds ratio of the enrichments are represented by color and dot size, respectively

Marker gene profiles (**Supplementary Figure S4B**) and gene ontology terms enriched in subtype-specific markers (**Figure 4C and Supplementary Figure S4C**) revealed that Macs-2 and Macs-3 displayed pronounced pro-inflammatory characteristics and were enriched for interferon-response signatures (**Supplementary Tables 2A and 2B**). Notably, Ifnb1 expression was highest in Macs-2 and was induced more rapidly following WD feeding compared to other macrophage subtypes (**Supplementary Figure S4B,D**), suggesting that this subtype may represent one potential local source of interferon signaling within islets during WD feeding. Macs-2 also exhibited a strong TNF- and NF-κB-driven transcriptional program, which was less prominent in other macrophage subtypes. In contrast, Macs-5 was characterized by high expression of phagocytic genes such as Ctsd, Prdx1, and Lgals3 (**Supplementary Figure S4B**), suggesting a phagocytic-like macrophage state.

The ratio of macrophage populations among the non-B-cell immune cell population increased one week after WD feeding, particularly the type I interferon-responsive macrophage subpopulation Macs-3 (**Figure 4C**). This observation suggested enhanced interferon-response programs within islet macrophages. To further investigate how Macs-3 respond to pro-inflammatory conditions, we conducted differential gene expression analyses. As predicted, both overnutrition and aging induced genes (e.g. *Cxcl9*, *Cxcl10*) [33] associated with a type I interferon response in Macs-3 (**Figure 4E-G; Supplementary Table 2C**). Interestingly, aging elicited a more canonical type I interferon response characterized by upregulation of *Stat1* and *Cxcl9* (**Figure 4G**) [33]. In contrast, WD feeding induced a distinct type I interferon response with elevated expression of *Stat3*-associated genes, including *Fos* and *Il1b* [34], alongside increased levels of the negative feedback regulator *Socs3* [33] (**Figure 4E,F**).

To further characterize the inflammatory responses, we categorized genes regulated in Macs-3 in response to overnutrition or aging into modules with shared expression dynamics across conditions (**Figure 4H**, **Supplementary Table 2D**). This analysis identified gene modules specific to aging, WD feeding, or shared between both conditions. Gene ontology and pathway enrichment analyses (**Figure 4I, Supplementary Table 2E**) revealed downregulation of genes associated with glycolytic processes in both WD feeding and aging (Module-1), suggesting metabolic reprogramming of Macs-3 under both conditions. Additionally, genes involved in cytoskeletal organization and RNA splicing were similarly downregulated. Conversely, genes upregulated in both WD feeding and aging (Module-7) were enriched for pathways related to cell proliferation, consistent with the expansion of Macs-3 within islets. Aging was associated with induction of a more canonical type I interferon-response program (Module-9, **Figure 4I**) together with suppression of the PERK pathway (Module-3, **Figure 4I**), a pathway previously implicated in immunosuppressive macrophage functions [35]. Genes specifically upregulated during WD feeding (Module-4) were enriched for mTORC1 signaling (**Figure 4I**), consistent with a non-canonical interferon-response program [36].

While these findings characterize the interferon response in Macs-3, we next sought to examine whether similar inflammatory programs were also present in the other major activated macrophage subtype, Macs-2. During overnutrition, this response involved STAT3 signaling, as evidenced by increased Socs3 expression one week after WD initiation (**Supplementary Figure S4E, Supplementary Table 2F**). WD feeding also triggered sustained expression of multiple inflammatory cytokine and chemokine genes in Macs-2 (**Supplementary Figure S4E,F, Supplementary Table 2F**), including Ifnb1, suggesting that this population may contribute to the local interferon signaling environment within islets. However, Macs-2 did not exhibit strong cytokine induction during aging (**Supplementary Figure S4G, Supplementary Table 2F**).

### Distinct responses of beta cells to overnutrition and aging

To investigate how pancreatic beta cells respond to overnutrition and aging, we first assessed the metabolic phenotypes of mice subjected to WD feeding or aging. Consistent with previous studies [37, 38], WD feeding rapidly altered insulin secretion and, by 12 weeks, caused glucose intolerance, insulin resistance, and compensatory hypersecretion (**Supplementary Figure S5A–H**). In contrast, aged mice maintained normal glucose tolerance and insulin sensitivity while exhibiting increased basal and glucose-stimulated insulin secretion (**Supplementary Figure S5I–L**). To characterize beta-cell transcriptional responses, we analyzed scRNA-seq data and identified three beta-cell subtypes by subclustering. These subclusters corresponded to previously described [39] mature (Beta-1 and Beta-2) and immature (Beta-3) β-cell states (**Figure 5A; Supplementary Figure S5M,N**). Among the mature populations, Beta-1 cells exhibited enhanced mTORC1 signaling, whereas Beta-2 cells were enriched for peptide metabolism pathways (**Figure 5B–D; Supplementary Tables 3A,B**). The relative abundance of beta-cell subtypes was unchanged by WD feeding or aging (**Figure 5E**).

**Figure 5:**
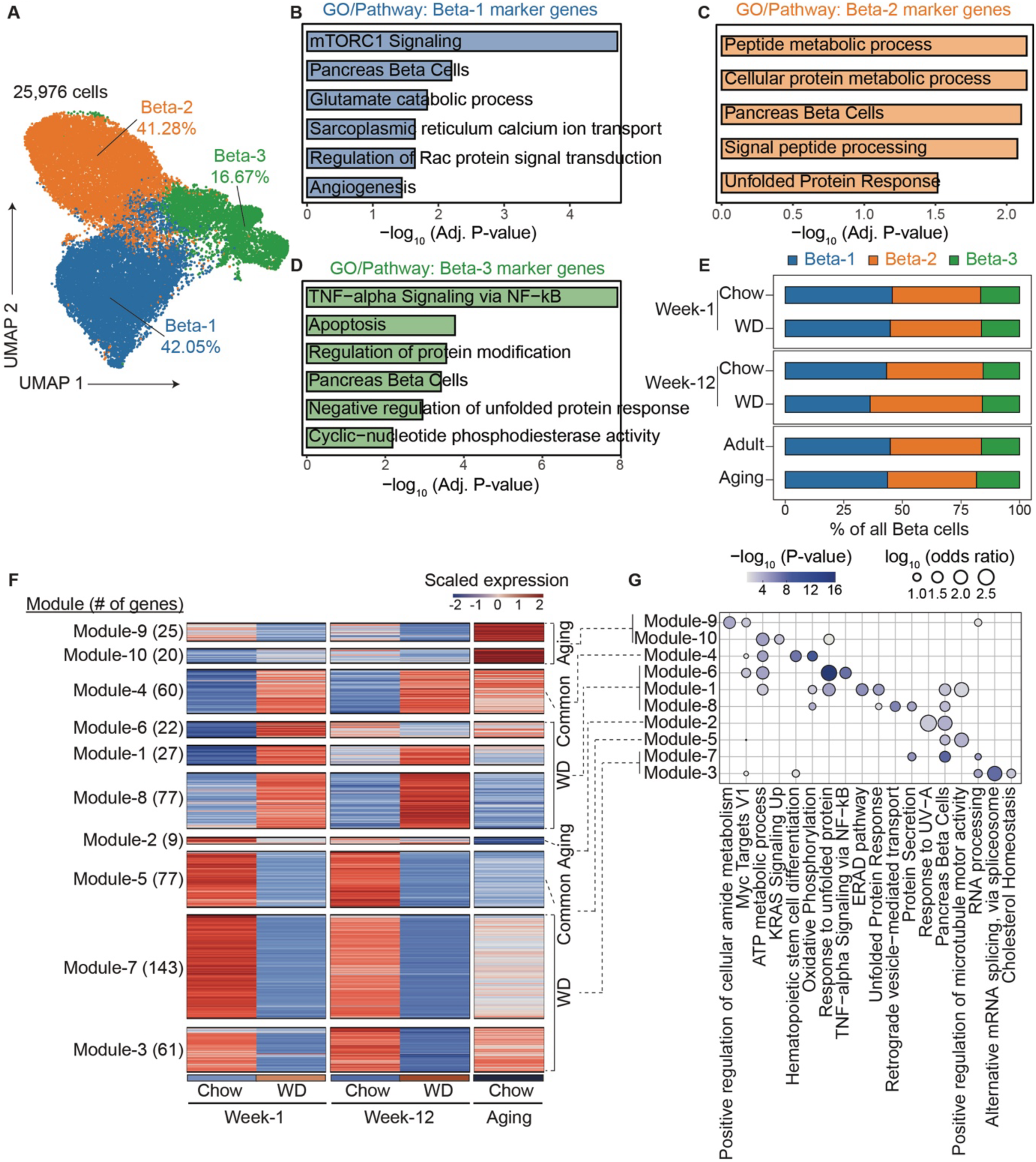
Western diet feeding and aging elicit distinct and time-dependent transcriptional changes in beta cells. **(A)** UMAP embedding of islet beta cells pooled across time points (week-1, week-12, and aging) and experimental conditions (WD and chow). The overall proportion of each defined subtype is indicated. **(B-D)** Enriched gene ontology (GO) terms for marker genes in Beta-1 **(B)**, Beta-2 **(C)** and Beta-3 **(D)**. Significance (-log_10_ adjusted p-value) of the enrichments is shown. **(E)** Proportions of beta cell subtypes expressed as percentages of all beta cells in each experimental group. Data were pooled across replicates for each group. **(F)** K-means clustering of all differentially expressed genes in beta cells across the five experimental groups. The color bar represents the scaled gene expression levels across all conditions analyzed. **(G)** Dot plot showing enriched gene ontology (GO) terms for selected k-means gene modules from **(F)**. Significance (-log_10_ p-value) and log_10_ odds ratio of the enrichments are represented by color and dot size, respectively.

Despite stable beta cell subtype composition, significant transcriptomic alterations occurred across all subtypes in response to overnutrition and aging, with similar direction and magnitude of changes (**Supplementary Figure S5O**). Given the overall consistency of transcriptional responses across beta-cell subtypes, we combined all beta cells for differential expression analysis, which identified gene modules that were specific to aging, specific to WD feeding, or shared between both conditions (**Figure 5F; Supplementary Table 3C**). Pathway analysis revealed that both WD feeding and aging enhanced oxidative phosphorylation while reducing beta-cell identity programs (**Figure 5G, Supplementary Table 3D**). WD feeding uniquely activated the unfolded protein response and suppressed RNA processing and splicing pathways, whereas aging-associated changes were more modest and included increased G6pc2 expression (**Supplementary Figure S5P**), consistent with enhanced glucose sensitivity and elevated basal insulin secretion (**Supplementary Figure S5L**). WD feeding also elicited temporally distinct responses, with an early TNF-α transcriptional program followed by enrichment of retrograde vesicle transport pathways during chronic feeding (**Figure 5F**). Notably, beta cells exhibited a TNF-α response at week 1 of WD feeding, aligning with increased Tnf expression in Macs-2 during the same period (**Supplementary Figure S4E**).

### Inferred communication between interferon-responsive macrophages and CD8⁺ cytotoxic T cells

To characterize T cells in the islet, we sub-clustered the scRNA-seq data (**Figure 3C**) and identified several T cell subpopulations, along with other lymphocytes such as innate lymphoid cells (ILC2/3), γδ T cells, and NK cells (**Figure 6A**). Both CD4⁺ and CD8⁺ T cells were annotated in our analysis, allowing the identification of several rare subpopulations. Within the CD4⁺ compartment, we identified two CD4^+^ naive T cell subtypes (naive-CD-1 and naive-CD4-2), Th1-like cells, T follicular helper (Tfh) cells and regulatory T cells (Tregs). Key CD8⁺ T cell subsets included CD8^+^ naive T cells, cytotoxic T cells (Eomes⁺), memory T cells (Il7r/CD127⁺), interferon-responsive cells (IFN responsive), and a proliferating CD8⁺ population (**Figure 6B, Supplementary Table 4A**). CD8^+^ cytotoxic T cells expressed classical cytotoxic T cell markers such as *Gzmk* (encoding granzyme) and *Ccl5* (**Figure 6B**). Although our single-cell analysis does not quantify the absolute number of lymphocytes within islets, we observed a marked enrichment of several T cell populations, most notably CD8⁺ cytotoxic T cells, among all CD19^-^ immune cell population in aged islets (**Figure 6C**).

**Figure 6:**
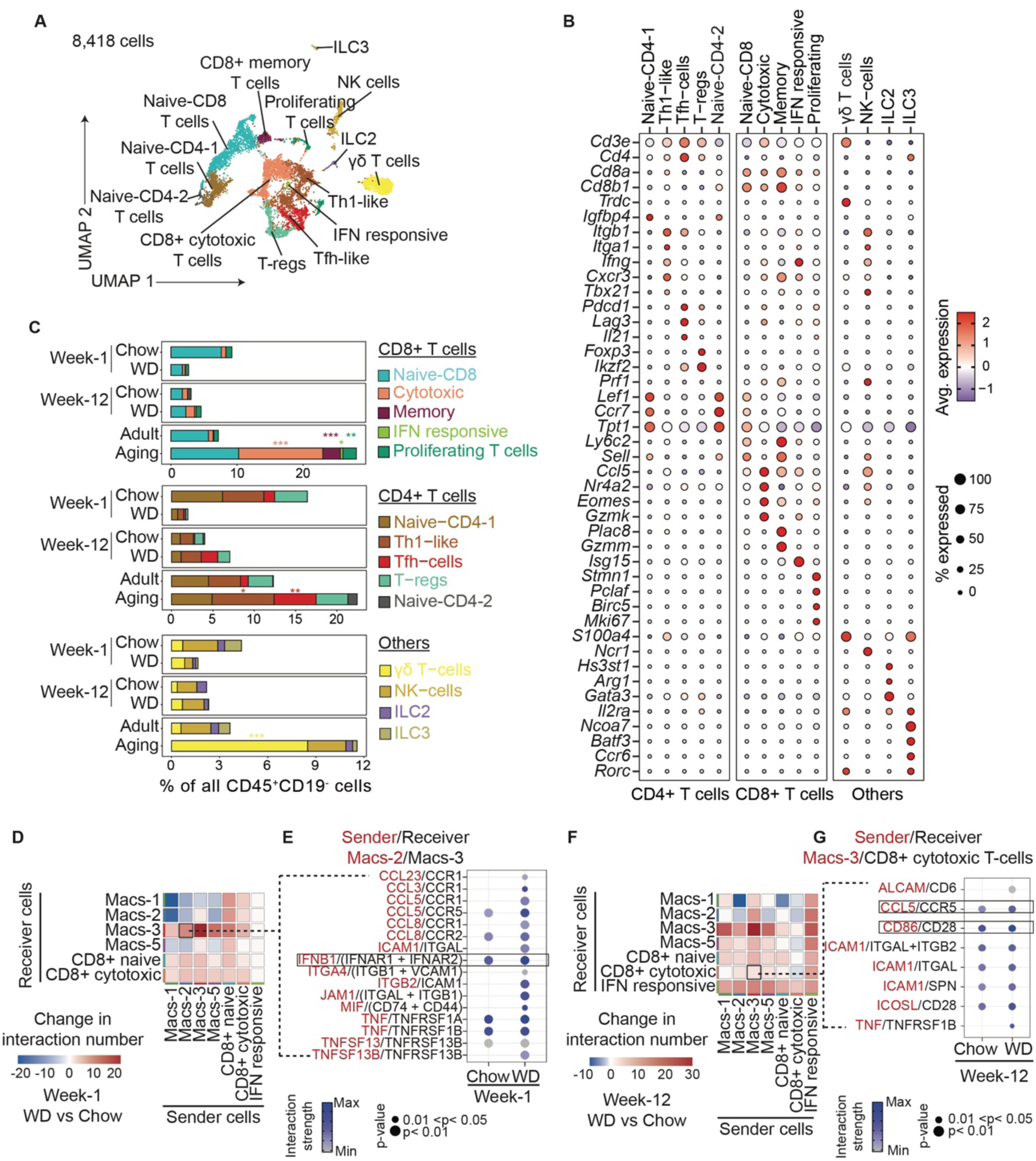
Western diet feeding leads to accumulation of CD8^+^ cytotoxic T cells in the pancreas. **(A)** scRNA-seq UMAP embedding of islet-associated T cells and other lymphocyte populations pooled across time points (week-1, week-12, and aging) and experimental conditions (WD and chow). Cell types were defined based on marker gene expression. **(B)** Dot plot of hallmark genes for cell types identified in **(A)**. **(C)** Proportions of CD8^+^ T cell, CD4^+^ T cells and other lymphocyte subpopulations expressed as percentages of all immune cell types, excluding B cells, across experimental groups. Data were pooled across conditions and replicates (see **Supplementary Figure S3A)**. * p<0.05, ** p<0.01, *** p < 0.001, calculated using a mixed-effects binomial model. **(D)** Heatmap showing changes in the number of ligand-receptor pairs among indicated cell populations, comparing week-1 chow and week-1 WD conditions (n=2 chohorts for WD, n=3 cohorts for chow). **(E)** Dot plots showing the strength of ligand-receptor interactions from Macs-2 to Macs-3 under week-1 chow and week-1 WD conditions. Highlighted are signals with increased interaction strength in the WD condition. Dot color and size represent the communication probability and p-values, calculated using a one-sided permutation test in CellChat. **(F)** Heatmap showing changes in the number of ligand-receptor pairs among indicated cell populations, comparing week-12 chow and week-12 WD conditions (n=2 cohorts for WD, n=3 cohorts for chow). **(G)** Dot plots showing the strength of ligand-receptor interactions from Macs-3 to CD8^+^ cytotoxic T cells under week-12 chow and week-12 WD conditions. Highlighted are signals with increased interaction strength in the WD condition. Dot color and size represent the communication probability and p-values, calculated using a one-sided permutation test in CellChat.

Recognizing that activated macrophages can stimulate T cells, we analyzed receptor-ligand interactions predicted from the scRNA-seq data (**see Methods**). Both acute (**Figure 6D,E, Supplementary Figure S6A, Supplementary Table 4B**) and chronic (**Figure 6F,G, Supplementary Figure S6A, Supplementary Table 4C**) overnutrition as well as aging (**Supplementary Figure S6A,B, Supplementary Table 4D**) altered intercellular communication among immune cells. One week into WD feeding, Macs-2 macrophages showed increased signaling to Macs-3 via interferon β1 and its receptors (**Figure 6E**), consistent with their role as the major source of interferon β1 (**Supplementary Figure S4B,D**) and with the heightened type I interferon response detected in Macs-3 (**Figure 4E**). Macs-3 also exhibited enhanced predicted communication with CD8^+^ cytotoxic T cells at week 12 of WD feeding via multiple strengthened ligand-receptor interactions (**Figure 6F**). Key interactions included Ccl5, secreted by Macs-3, binding to the Ccr5 receptor on CD8^+^ cytotoxic T cells, and co-stimulatory signaling via Cd86 from Macs-3 and the Cd28 receptor on T cells (**Figure 6G**).

Interestingly, we observed increased interactions among all examined immune cell subpopulations in the aging cohort compared to adults, suggesting an altered landscape of immune crosstalk during aging (**Supplementary Figure S6B**). Notably, in the aging cohort, enriched interactions between both Macs-2 and Macs-3 with CD8⁺ cytotoxic T cells did not involve Ccl5. Instead, they featured Ccl3 and Ccl4 binding to the Ccr5 receptor, and Ccl8 binding to the Ccr2 receptor on CD8⁺ cytotoxic T cells (**Supplementary Figure S6C,D**). Despite these differences between aging and WD-induced interactions, the Cd86–Cd28 co-stimulatory interaction between Macs-3 and CD8⁺ cytotoxic T cells was consistently shared across both metabolic stress conditions (**Supplementary Figure S6D**). These findings reveal additional distinctions in the inflammatory landscape within islets between WD feeding and aging, including differences in predicted macrophage–T cell communication pathways.

## DISCUSSION

We combined imaging mass cytometry (IMC) and single-cell RNA sequencing (scRNA-seq) to define the composition, spatial organization, and transcriptional states of pancreatic immune cells during overnutrition and aging.

IMC enabled quantification of the absolute abundance, density, and spatial distribution of immune cell populations throughout the pancreas. Using this approach, we found that the F4/80^low^ macrophage subtype accumulated under both WD feeding and aging, with expansion occurring predominantly in the exocrine compartment during WD feeding but extending to both the exocrine and peri-islet regions in aged mice. IMC also revealed widespread accumulation of activated effector-like CD8^+^ T cells throughout the pancreas during WD feeding, whereas this response was not observed during aging. In contrast, scRNA-seq provided high-resolution characterization of immune cell composition and transcriptional states within pancreatic islets. This analysis identified an increased relative abundance of the type I interferon-responsive Macs-3 macrophage subtype following one week of WD feeding and enrichment of CD8^+^ cytotoxic T cells in aged islets. Because scRNA-seq was performed on FACS-enriched CD45^+^ cells, it assessed the relative composition of islet immune populations rather than their absolute abundance. Together, these complementary approaches reveal both compartment-specific changes in immune cell distribution and condition-specific remodeling of islet immune cell states.

Despite this limitation, our scRNA-seq profiling of FACS-enriched islet CD45^+^ cells provided high-resolution insight into immune cell heterogeneity and its molecular remodeling under stress conditions. Our analysis identified five macrophage subtypes within the islet, each with distinct activation states. Among them, two subtypes characterized by relatively low F4/80 expression (Macs-2 and Macs-3) exhibited the strongest activation signatures. The Macs-3 subtype showed robust type I interferon responsiveness, whereas the Macs-2 subtype emerged as the primary source of type I interferon production. Previous studies have shown preferential expansion of F4/80low macrophages in obese islets [14, 17]. Our data extend these observations by resolving two activated F4/80low macrophage subtypes with distinct functions, identifying Macs-3 as the expanding interferon-responsive population and Macs-2 as a putative signaling population.

In addition, the mechanism of macrophage expansion in islets has been debated, with uncertainty regarding whether it arises from monocyte infiltration or local proliferation [13]. Our identification of a proliferating macrophage subtype (Macs-4), together with the absence of peri-islet macrophage accumulation by IMC, supports local proliferation as a contributor to macrophage expansion during overnutrition.

Chronic inflammation associated with overnutrition (“metaflammation”) and aging (“inflammaging”) share common features, including activation of innate immune compartments, particularly macrophages [4, 40, 41]. Although these inflammatory states have been compared extensively in systemic metabolism and adipose tissue [4], their relationship within the pancreas and pancreatic islets remains poorly understood. Our integrated IMC and scRNA-seq analyses provide a framework for comparing pancreatic immune remodeling during metabolic stress and aging, two major risk factors for type 2 diabetes. We identified common signatures of inflammation under both aging and overnutrition, including the accumulation of a F4/80^low^ macrophage population throughout the pancreas and a heightened type I interferon response within the islet-associated Macs-3 subtype. These findings are consistent with the concept that metaflammation accelerates inflammaging [42].

Transcriptome-wide analysis further revealed that the nature of the type I interferon response in Macs-3 differed between the two conditions. Aging was associated with a more canonical type I interferon-response signature, whereas overnutrition elicited a broader response that included STAT3-associated transcriptional programs and mTORC1 signaling. Given that STAT3 suppresses type I interferon activity [33], these observations may reflect an intrinsic mechanism that islet-associated macrophages utilize to temper inflammatory signaling during overnutrition. Age-related disruption of this regulatory network could influence pancreatic inflammatory responses and thereby contribute to the pathogenesis of type 2 diabetes.

T-cell accumulation has been reported in both the exocrine pancreas and pancreatic islets of patients with type 2 diabetes [10, 17, 43, 44]. In rodents, increased T-cell abundance has been observed in islets of aged mice [23], but not in animals subjected to high-fat diet feeding [17]. Our scRNA-seq analysis was consistent with these observations within islets, whereas IMC revealed widespread accumulation of activated effector-like CD8^+^ T cells throughout the pancreas during WD feeding but not aging. These findings suggest that metabolic stress and aging elicit distinct compartment-specific T-cell responses. Receptor-ligand analysis further identified enhanced communication between islet macrophages and CD8^+^ cytotoxic T cells under both conditions, although the functional significance of this interaction remains to be determined.

In summary, our integrated spatial and single-cell analyses demonstrate that metabolic stress and aging remodel pancreatic inflammation through overlapping yet distinct immune programs. By resolving immune cell composition, spatial organization, and transcriptional states across the pancreas and pancreatic islets, this study provides a framework for understanding how immune heterogeneity contributes to pancreatic dysfunction in metabolic disease and aging.

## ACKNOWLEDGMENTS

The authors thank all members of the participating laboratories for helpful discussions and technical assistance.

## Author Contributions

S.S. and I.O. contributed equally to this work and performed the majority of experiments and analyses. S.S., I.O., M.S., B.S., and H.Z. conceived and designed the study. S.S., I.O., K.M., M.Sch., F.L., S.Sr., J.C.J., T.T., and H.Z. acquired data. S.S., I.O., M.B., T.C., T.B., B.S., M.S., and H.Z. analyzed and interpreted the data. S.S., I.O., and H.Z. drafted the manuscript. All authors contributed to data interpretation, critically revised the manuscript for important intellectual content, and approved the final version of the manuscript.

## Guarantor Statement

H.Z. is the guarantor of this work and, as such, had full access to all the data in the study and takes responsibility for the integrity of the data and the accuracy of the data analysis.

## Prior Presentation

Parts of this study have not been previously presented in abstract or meeting form.

## Artificial Intelligence Disclosure

During the course of preparing this work, the authors used ChatGPT (OpenAI) to assist with language editing and improving the clarity of the manuscript. Following the use of this tool, the authors reviewed all content for scientific accuracy, revised the text as necessary, and take full responsibility for all content of this publication.

## REFERENCES

1. Khan MAB, Hashim MJ, King JK, Govender RD, Mustafa H, Al Kaabi J (2020) Epidemiology of Type 2 Diabetes - Global Burden of Disease and Forecasted Trends. J Epidemiol Glob Health 10(1):107–111. 10.2991/jegh.k.191028.001

2. Schulze MB, Hoffmann K, Manson JE, et al (2005) Dietary pattern, inflammation, and incidence of type 2 diabetes in women. Am J Clin Nutr 82(3):675–684; quiz 714–715. 10.1093/ajcn.82.3.675

3. Donath MY, Dalmas É, Sauter NS, Böni-Schnetzler M (2013) Inflammation in Obesity and Diabetes: Islet Dysfunction and Therapeutic Opportunity. Cell Metabolism 17(6):860–872. 10.1016/j.cmet.2013.05.001

4. Prattichizzo F, De Nigris V, Spiga R, et al (2018) Inflammageing and metaflammation: The yin and yang of type 2 diabetes. Ageing Research Reviews 41:1–17. 10.1016/j.arr.2017.10.003

5. Lee YS, Wollam J, Olefsky JM (2018) An Integrated View of Immunometabolism. Cell 172(1):22–40. 10.1016/j.cell.2017.12.025

6. Ying W, Fu W, Lee YS, Olefsky JM (2020) The role of macrophages in obesity-associated islet inflammation and β-cell abnormalities. Nat Rev Endocrinol 16(2):81–90. 10.1038/s41574-019-0286-3

7. Gregor MF, Hotamisligil GS (2011) Inflammatory mechanisms in obesity. Annu Rev Immunol 29:415–445. 10.1146/annurev-immunol-031210-101322

8. Hotamisligil GS (2017) Inflammation, metaflammation and immunometabolic disorders. Nature 542(7640):177–185. 10.1038/nature21363

9. Böni-Schnetzler M, Meier DT (2019) Islet inflammation in type 2 diabetes. Semin Immunopathol 41(4):501–513. 10.1007/s00281-019-00745-4

10. Ehses JA, Perren A, Eppler E, et al (2007) Increased number of islet-associated macrophages in type 2 diabetes. Diabetes 56(9):2356–2370. 10.2337/db06-1650

11. Böni-Schnetzler M, Thorne J, Parnaud G, et al (2008) Increased interleukin (IL)-1beta messenger ribonucleic acid expression in beta -cells of individuals with type 2 diabetes and regulation of IL-1beta in human islets by glucose and autostimulation. J Clin Endocrinol Metab 93(10):4065–4074. 10.1210/jc.2008-0396

12. Eguchi K, Nagai R (2017) Islet inflammation in type 2 diabetes and physiology. J Clin Invest 127(1):14–23. 10.1172/JCI88877

13. Ying W, Fu W, Lee YS, Olefsky JM (2020) The role of macrophages in obesity-associated islet inflammation and β-cell abnormalities. Nat Rev Endocrinol 16(2):81–90. 10.1038/s41574-019-0286-3

14. Cucak H, Grunnet LG, Rosendahl A (2014) Accumulation of M1-like macrophages in type 2 diabetic islets is followed by a systemic shift in macrophage polarization. J Leukoc Biol 95(1):149–160. 10.1189/jlb.0213075

15. Eguchi K, Manabe I, Oishi-Tanaka Y, et al (2012) Saturated fatty acid and TLR signaling link β cell dysfunction and islet inflammation. Cell Metab 15(4):518–533. 10.1016/j.cmet.2012.01.023

16. Westwell-Roper CY, Ehses JA, Verchere CB (2014) Resident macrophages mediate islet amyloid polypeptide-induced islet IL-1β production and β-cell dysfunction. Diabetes 63(5):1698–1711. 10.2337/db13-0863

17. Ying W, Lee YS, Dong Y, et al (2019) Expansion of Islet-Resident Macrophages Leads to Inflammation Affecting β Cell Proliferation and Function in Obesity. Cell Metab 29(2):457–474.e5. 10.1016/j.cmet.2018.12.003

18. Rohm TV, Meier DT, Olefsky JM, Donath MY (2022) Inflammation in obesity, diabetes, and related disorders. Immunity 55(1):31–55. 10.1016/j.immuni.2021.12.013

19. Kiran S, Kumar V, Murphy EA, Enos RT, Singh UP (2021) High Fat Diet-Induced CD8+ T Cells in Adipose Tissue Mediate Macrophages to Sustain Low-Grade Chronic Inflammation. Front Immunol 12:680944. 10.3389/fimmu.2021.680944

20. Janjuha S, Singh SP, Tsakmaki A, et al (2018) Age-related islet inflammation marks the proliferative decline of pancreatic beta-cells in zebrafish. Elife 7:e32965. 10.7554/eLife.32965

21. He W, Yuan T, Choezom D, et al (2018) Ageing potentiates diet-induced glucose intolerance, β-cell failure and tissue inflammation through TLR4. Sci Rep 8(1):2767. 10.1038/s41598-018-20909-w

22. Sandovici I, Hammerle CM, Cooper WN, et al (2016) Ageing is associated with molecular signatures of inflammation and type 2 diabetes in rat pancreatic islets. Diabetologia 59(3):502–511. 10.1007/s00125-015-3837-8

23. Denroche HC, Miard S, Sallé-Lefort S, Picard F, Verchere CB (2021) T cells accumulate in non-diabetic islets during ageing. Immunity & Ageing 18(1):8. 10.1186/s12979-021-00221-4

24. Ginhoux F, Yalin A, Dutertre CA, Amit I (2022) Single-cell immunology: Past, present, and future. Immunity 55(3):393–404. 10.1016/j.immuni.2022.02.006

25. Bressan D, Battistoni G, Hannon GJ (2023) The dawn of spatial omics. Science 381(6657):eabq4964. 10.1126/science.abq4964

26. Chiou J, Geusz RJ, Okino M-L, et al (2021) Interpreting type 1 diabetes risk with genetics and single-cell epigenomics. Nature 594(7863):398–402. 10.1038/s41586-021-03552-w

27. Damond N, Engler S, Zanotelli VRT, et al (2019) A Map of Human Type 1 Diabetes Progression by Imaging Mass Cytometry. Cell Metab 29(3):755–768.e5. 10.1016/j.cmet.2018.11.014

28. Wang YJ, Traum D, Schug J, et al (2019) Multiplexed In Situ Imaging Mass Cytometry Analysis of the Human Endocrine Pancreas and Immune System in Type 1 Diabetes. Cell Metab 29(3):769–783.e4. 10.1016/j.cmet.2019.01.003

29. Zakharov PN, Hu H, Wan X, Unanue ER (2020) Single-cell RNA sequencing of murine islets shows high cellular complexity at all stages of autoimmune diabetes. J Exp Med 217(6):e20192362. 10.1084/jem.20192362

30. Wang G, Chiou J, Zeng C, et al (2023) Integrating genetics with single-cell multiomic measurements across disease states identifies mechanisms of beta cell dysfunction in type 2 diabetes. Nat Genet 55(6):984–994. 10.1038/s41588-023-01397-9

31. Weng C, Gu A, Zhang S, et al (2023) Single cell multiomic analysis reveals diabetes-associated β-cell heterogeneity driven by HNF1A. Nat Commun 14(1):5400. 10.1038/s41467-023-41228-3

32. Levine JH, Simonds EF, Bendall SC, et al (2015) Data-Driven Phenotypic Dissection of AML Reveals Progenitor-like Cells that Correlate with Prognosis. Cell 162(1):184–197. 10.1016/j.cell.2015.05.047

33. Tsai M-H, Pai L-M, Lee C-K (2019) Fine-Tuning of Type I Interferon Response by STAT3. Front Immunol 10:1448. 10.3389/fimmu.2019.01448

34. Balic JJ, Albargy H, Luu K, et al (2020) STAT3 serine phosphorylation is required for TLR4 metabolic reprogramming and IL-1β expression. Nat Commun 11(1):3816. 10.1038/s41467-020-17669-5

35. Raines LN, Zhao H, Wang Y, et al (2022) PERK is a critical metabolic hub for immunosuppressive function in macrophages. Nat Immunol 23(3):431–445. 10.1038/s41590-022-01145-x

36. Mazewski C, Perez RE, Fish EN, Platanias LC (2020) Type I Interferon (IFN)-Regulated Activation of Canonical and Non-Canonical Signaling Pathways. Front Immunol 11:606456. 10.3389/fimmu.2020.606456

37. Ramms B, Pollow DP, Zhu H, et al (2022) Systemic LSD1 Inhibition Prevents Aberrant Remodeling of Metabolism in Obesity. Diabetes 71(12):2513–2529. 10.2337/db21-1131

38. De Leon ER, Brinkman JA, Fenske RJ, et al (2018) Age-Dependent Protection of Insulin Secretion in Diet Induced Obese Mice. Sci Rep 8(1):17814. 10.1038/s41598-018-36289-0

39. Sachs S, Bastidas-Ponce A, Tritschler S, et al (2020) Targeted pharmacological therapy restores β-cell function for diabetes remission. Nat Metab 2(2):192–209. 10.1038/s42255-020-0171-3

40. Franceschi C, Garagnani P, Parini P, Giuliani C, Santoro A (2018) Inflammaging: a new immune–metabolic viewpoint for age-related diseases. Nat Rev Endocrinol 14(10):576–590. 10.1038/s41574-018-0059-4

41. Qu L, Matz AJ, Karlinsey K, Cao Z, Vella AT, Zhou B (2022) Macrophages at the Crossroad of Meta-Inflammation and Inflammaging. Genes 13(11):2074. 10.3390/genes13112074

42. Franceschi C, Garagnani P, Morsiani C, et al (2018) The Continuum of Aging and Age-Related Diseases: Common Mechanisms but Different Rates. Front Med (Lausanne) 5:61. 10.3389/fmed.2018.00061

43. Butcher MJ, Hallinger D, Garcia E, et al (2014) Association of proinflammatory cytokines and islet resident leucocytes with islet dysfunction in type 2 diabetes. Diabetologia 57(3):491–501. 10.1007/s00125-013-3116-5

44. Wu M, Lee MYY, Bahl V, et al (2021) Single-cell analysis of the human pancreas in type 2 diabetes using multi-spectral imaging mass cytometry. Cell Reports 37(5):109919. 10.1016/j.celrep.2021.109919

