## Supplemental Figures for "Distinct macrophage and T cell programs shape pancreatic inflammation during metabolic stress and aging"

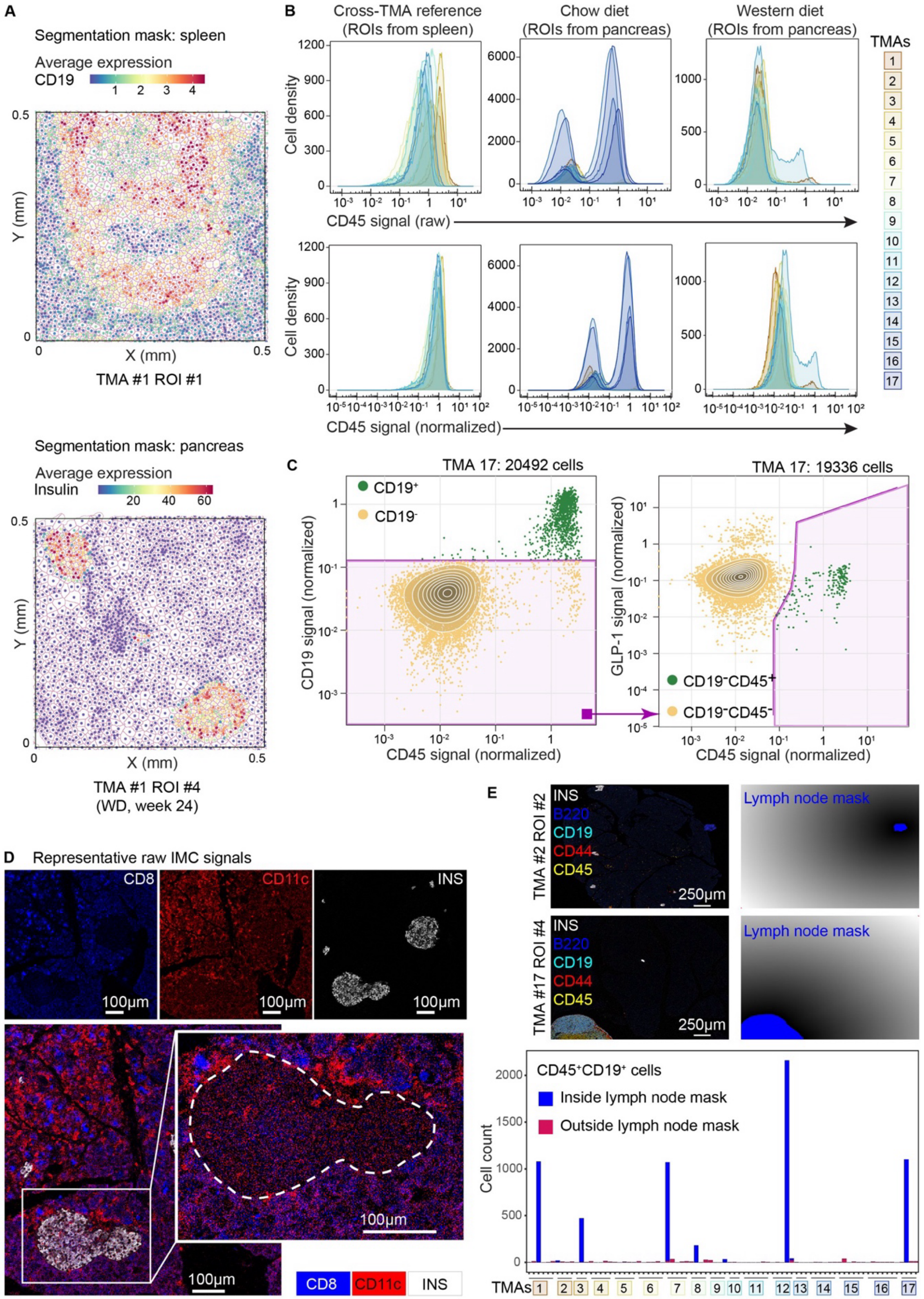

A
Segmentation mask: spleen
Average expression
CD19
Y (mm)
X (mm)
TMA #1 ROI #1
Segmentation mask: pancreas
Average expression
Insulin
TMA #1 ROI #4
(WD, week 24)
B
Cross-TMA reference
(ROIs from spleen)
Chow diet
(ROIs from pancreas)
Western diet
(ROIs from pancreas)
TMAs
Cell density
CD45 signal (raw)
CD45 signal (normalized)
C
TMA 17: 20492 cells
CD19+
CD19-
CD19 signal (normalized)
CD45 signal (normalized)
TMA 17: 19336 cells
GLP-1 signal (normalized)
CD19-CD45+
CD19-CD45-
D
Representative raw IMC signals
CD8
CD11c
INS
100μm
E
TMA #2 ROI #2
TMA #17 ROI #4
INS
B220
CD19
CD44
CD45
250μm
Lymph node mask
CD45+CD19+ cells
Inside lymph node mask
Outside lymph node mask
Cell count
TMAs

**Supplementary Figure 1: Quality control of IMC data.**

**(A)** Representative images showing cell segmentations and the expression of marker proteins on spleen (CD19, top) and pancreas (insulin, bottom) sections. The color of the dot at the center of each cell segmentation reflects the average expression of CD19 (top) and insulin (bottom). The dimensions of each region of interest (ROI) are indicated along both axes.

**(B)** Histograms showing raw (top) and reference-ROI-normalized (bottom) IMC signal intensity from the CD45 channel across all TMAs. ROIs from different TMAs are distinguished by varying colors.

**(C)** Representative scatter plots depicting the Boolean gating strategy employed to identify pancreatic non-B cell immune cells ($CD19^{-}CD45^{+}GLP1R^{-}$) within each TMA.

**(D)** Representative ROI showing raw signals from the CD8 (blue), CD11c (red), and insulin (white) channels. Scale bar, 100 µm.

**(E)** Representative ROIs showing two pancreatic lymph nodes (highlighted in blue). Pancreatic lymph node masks were generated based on a composite overlay of signals from B220, CD19, CD44, and CD45 (top right, see SI), scale bar, 250 µm. Bottom: Bar plot with counts of $CD19^{+}$ B cells within and outside of these lymph node masks.

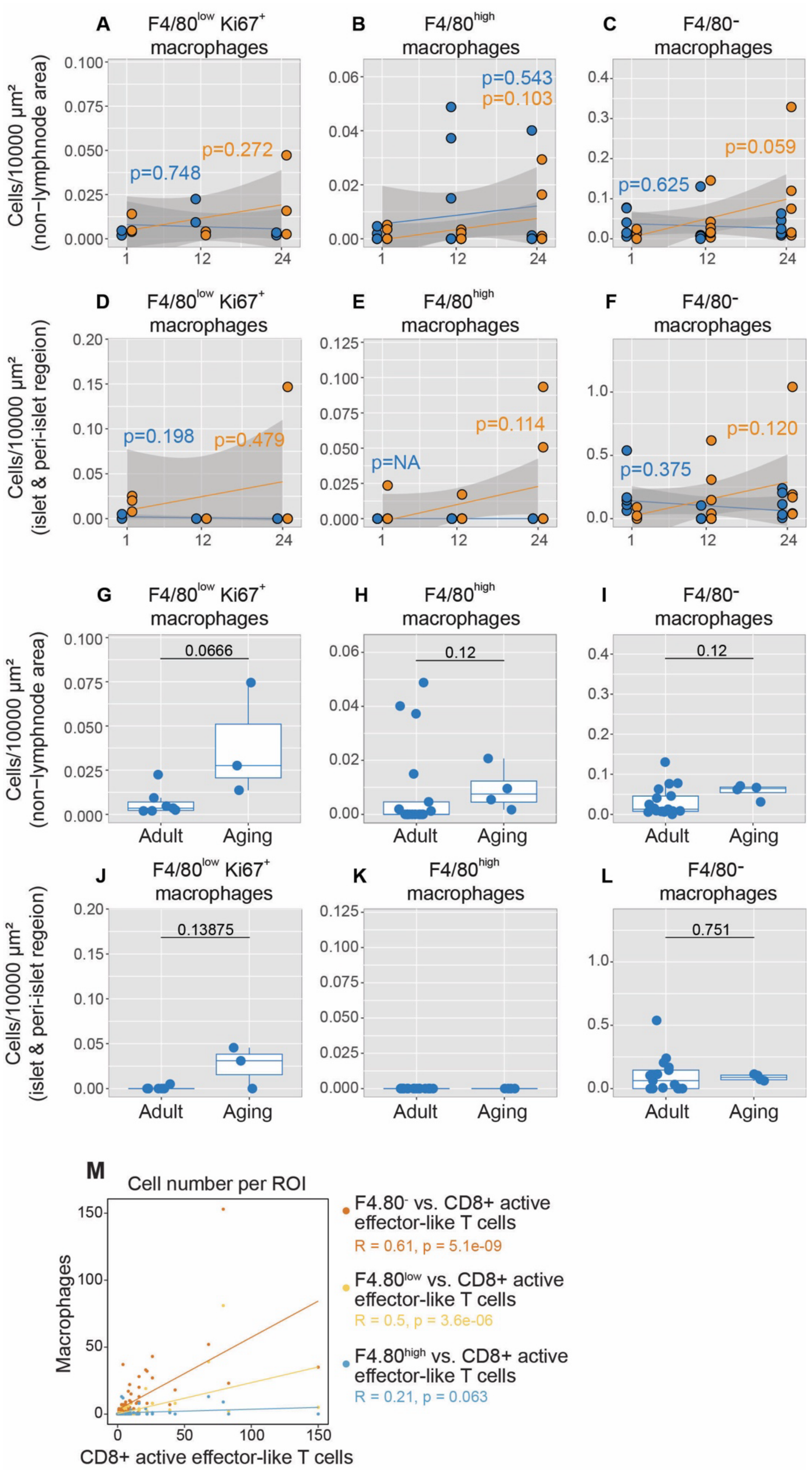
A
F4/80low Ki67+ macrophages
B
F4/80high macrophages
C
F4/80- macrophages
Cells/10000 µm² (non−lymphnode area)
p=0.748
p=0.272
p=0.543
p=0.103
p=0.625
p=0.059
D
F4/80low Ki67+ macrophages
E
F4/80high macrophages
F
F4/80- macrophages
Cells/10000 µm² (islet & peri-islet regeion)
p=0.198
p=0.479
p=NA
p=0.114
p=0.375
p=0.120
G
F4/80low Ki67+ macrophages
H
F4/80high macrophages
I
F4/80- macrophages
Cells/10000 µm² (non−lymphnode area)
0.0666
0.12
0.12
Adult
Aging
J
F4/80low Ki67+ macrophages
K
F4/80high macrophages
L
F4/80- macrophages
Cells/10000 µm² (islet & peri-islet regeion)
0.13875
0.751
Adult
Aging
M
Cell number per ROI
Macrophages
CD8+ active effector-like T cells
F4.80- vs. CD8+ active effector-like T cells
R = 0.61, p = 5.1e-09
F4.80low vs. CD8+ active effector-like T cells
R = 0.5, p = 3.6e-06
F4.80high vs. CD8+ active effector-like T cells
R = 0.21, p = 0.063

**Supplementary Figure 2: Cell density quantification and spatial correlation analysis.**

(**A-C**) Scatter plots depicting the cell counts of $F4/80^{low}$ $Ki67^{+}$ (**A**), $F4/80^{high}$ (**B**) and $F4/80^{-}$ (**C**) macrophages per 10000 $\mu m^2$of the pancreas (excluding lymph nodes) across different time points and feeding conditions. Solid lines represent linear regression fits, with shaded areas indicating 95% confidence intervals calculated from the standard error. Each condition includes data from $n \geq 3$ mice. P-values for the regression coefficient slope were determined using a linear model and t-tests.

(**D-F**) Scatter plots showing the cell counts of $F4/80^{low}$ $Ki67^{+}$ (**D**), $F4/80^{high}$ (**E**) and $F4/80^{-}$ (**F**) macrophages per 10000 $\mu m^2$ of the islets and peri-islet regions over time under feeding conditions. Linear regression fits (solid lines) and 95% confidence intervals (shaded areas) are displayed. Each condition includes data from $n \geq 3$ mice. P-values for the regression coefficient slope were determined using a linear model and t-tests.

(**G-I**) Box plots showing the cell counts of $F4/80^{low}$ $Ki67^{+}$ (**G**), $F4/80^{high}$ (**H**) and $F4/80^{-}$ (**I**) macrophages per 10000 $\mu m^2$ of the pancreas (excluding lymph nodes) in adult and aging mice. Each condition includes data from $n \geq 3$ mice. P-values were determined by Wilcoxon signed-rank test with Benjamini & Hochberg adjustment.

(**J-L**) Box plots showing the cell counts of $F4/80^{low}$ $Ki67^{+}$ (**J**), $F4/80^{high}$ (**K**) and $F4/80^{-}$ (**L**) macrophages per 10000 $\mu m^2$ of islets and peri-islet regions in adult and aging mice. Each condition includes data from $n \geq 3$ mice. P-values were determined by Wilcoxon signed-rank test with Benjamini & Hochberg adjustment.

(**M**) Scatter plot showing the Pearson correlation between the counts of CD8+ activated effector-like T cells and three macrophage subpopulations ($F4/80^{-}$, $F4/80^{low}$ and $F4/80^{high}$) within the same regions of interest (ROIs). Solid lines indicate the best fit for each correlation analysis, with Pearson correlation coefficients and p-values provided for the corresponding macrophage subsets.

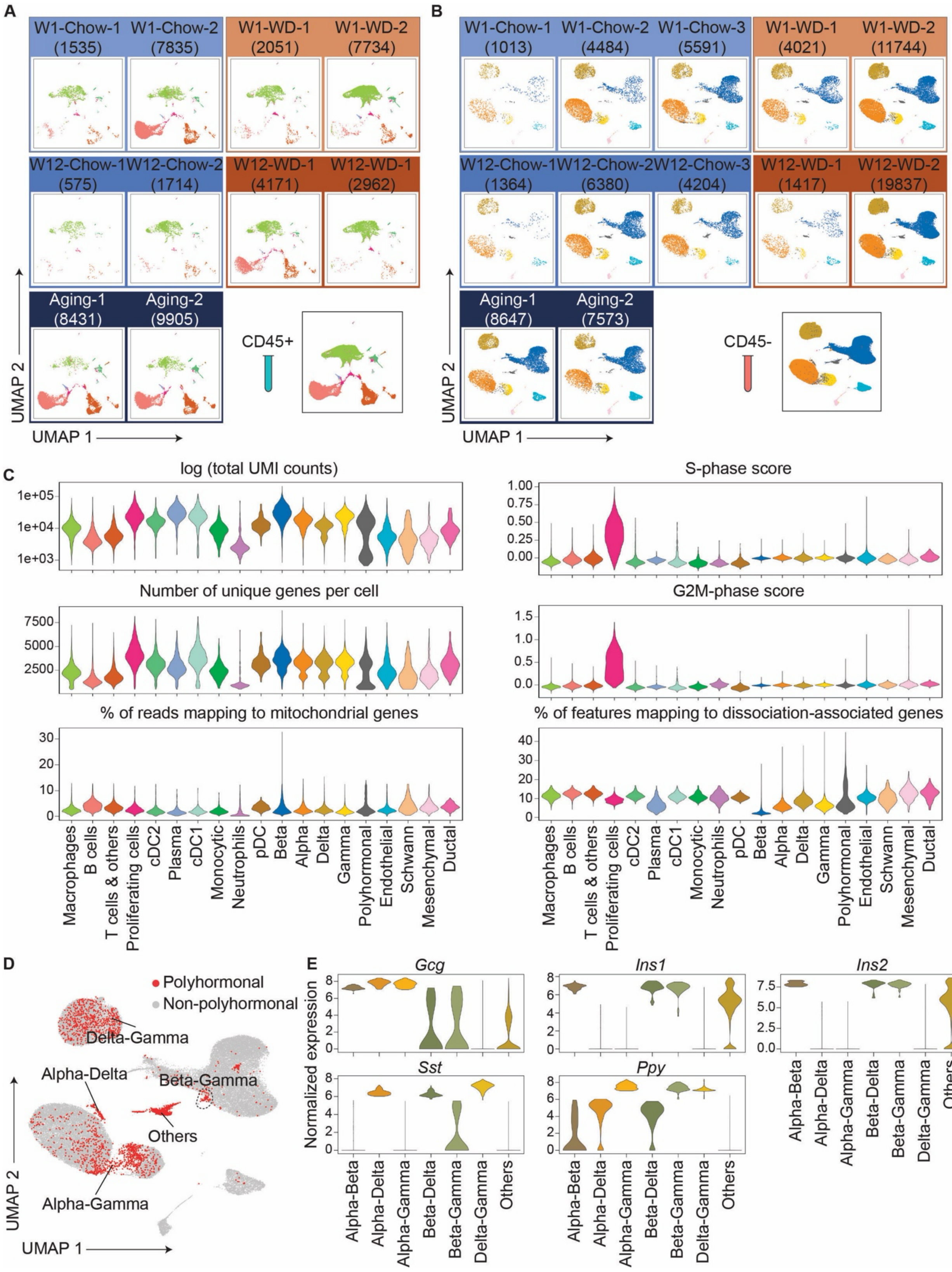

A
W1-Chow-1 (1535)
W1-Chow-2 (7835)
W1-WD-1 (2051)
W1-WD-2 (7734)
W12-Chow-1 (575)
W12-Chow-2 (1714)
W12-WD-1 (4171)
W12-WD-1 (2962)
Aging-1 (8431)
Aging-2 (9905)
CD45+
UMAP 2
UMAP 1
B
W1-Chow-1 (1013)
W1-Chow-2 (4484)
W1-Chow-3 (5591)
W1-WD-1 (4021)
W1-WD-2 (11744)
W12-Chow-1 (1364)
W12-Chow-2 (6380)
W12-Chow-3 (4204)
W12-WD-1 (1417)
W12-WD-2 (19837)
Aging-1 (8647)
Aging-2 (7573)
CD45-
UMAP 2
UMAP 1
C
log (total UMI counts)
Number of unique genes per cell
% of reads mapping to mitochondrial genes
S-phase score
G2M-phase score
% of features mapping to dissociation-associated genes
Macrophages
B cells
T cells & others
Proliferating cells
cDC2
Plasma
cDC1
Monocytic
Neutrophils
pDC
Beta
Alpha
Delta
Gamma
Polyhormonal
Endothelial
Schwann
Mesenchymal
Ductal
D
Polyhormonal
Non-polyhormonal
Delta-Gamma
Alpha-Delta
Beta-Gamma
Others
Alpha-Gamma
UMAP 2
UMAP 1
E
Gcg
Ins1
Ins2
Sst
Ppy
Normalized expression
Alpha-Beta
Alpha-Delta
Alpha-Gamma
Beta-Delta
Beta-Gamma
Delta-Gamma
Others

**Supplementary Figure 3: Quality control of scRNA-seq data.**

**(A** and **B)** Split UMAPs depicting the distribution of the annotated immune **(A**, $CD45^{+}$**)** and non-immune **(B,** $CD45^{-}$**)** cell types across experimental cohorts.

**(C)** Violin plots depicting quality control metrics for scRNA-seq data. In addition to standard quality control metrics, we also examined genes induced by enzymatic dissociation (**Supplementary Table 1D**).

**(D)** UMAP embedding of non-immune cells in grey. Polyhormonal cells are highlighted in red.

**(E)** Violin plots depicting the expression of five endocrine hormone genes across annotated polyhormonal cells.

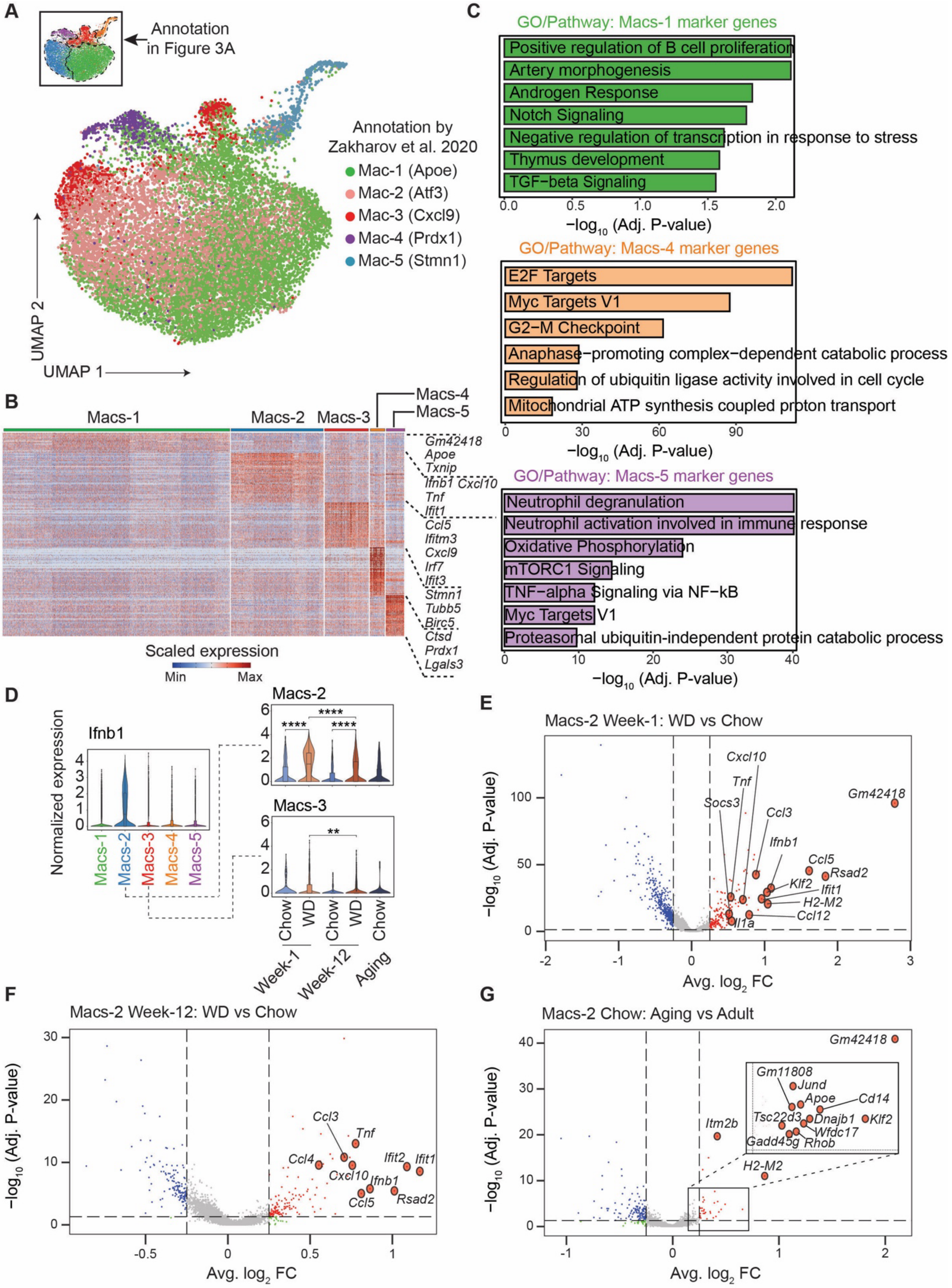

A
Annotation in Figure 3A
Annotation by Zakharov et al. 2020
Mac-1 (Apoe)
Mac-2 (Atf3)
Mac-3 (Cxcl9)
Mac-4 (Prdx1)
Mac-5 (Stmn1)
UMAP 2
UMAP 1
B
Macs-1
Macs-2
Macs-3
Macs-4
Macs-5
Gm42418
Apoe
Txnip
Ifnb1 Cxcl10
Tnf
Ifit1
Ccl5
Ifitm3
Cxcl9
Irf7
Ifit3
Stmn1
Tubb5
Birc5
Ctsd
Prdx1
Lgals3
Scaled expression
Min
Max
C
GO/Pathway: Macs-1 marker genes
Positive regulation of B cell proliferation
Artery morphogenesis
Androgen Response
Notch Signaling
Negative regulation of transcription in response to stress
Thymus development
TGF−beta Signaling
−log10 (Adj. P-value)
GO/Pathway: Macs-4 marker genes
E2F Targets
Myc Targets V1
G2−M Checkpoint
Anaphase−promoting complex−dependent catabolic process
Regulation of ubiquitin ligase activity involved in cell cycle
Mitochondrial ATP synthesis coupled proton transport
−log10 (Adj. P-value)
GO/Pathway: Macs-5 marker genes
Neutrophil degranulation
Neutrophil activation involved in immune response
Oxidative Phosphorylation
mTORC1 Signaling
TNF−alpha Signaling via NF−kB
Myc Targets V1
Proteasomal ubiquitin-independent protein catabolic process
−log10 (Adj. P-value)
D
Ifnb1
Normalized expression
Macs-1
Macs-2
Macs-3
Macs-4
Macs-5
Macs-2
Macs-3
Chow
WD
Chow
WD
Chow
Week-1
Week-12
Aging
E
Macs-2 Week-1: WD vs Chow
Cxcl10
Tnf
Socs3
Ccl3
Gm42418
Ifnb1
Ccl5
Rsad2
Klf2
Ifit1
H2-M2
Ccl12
Il1a
−log10 (Adj. P-value)
Avg. log2 FC
F
Macs-2 Week-12: WD vs Chow
Ccl3
Tnf
Ccl4
Cxcl10
Ifit2
Ifit1
Ifnb1
Ccl5
Rsad2
−log10 (Adj. P-value)
Avg. log2 FC
G
Macs-2 Chow: Aging vs Adult
Gm42418
Gm11808
Jund
Apoe
Cd14
Tsc22d3
Dnajb1
Klf2
Wfdc17
Gadd45g
Rhob
Itm2b
H2-M2
−log10 (Adj. P-value)
Avg. log2 FC

**Supplementary Figure 4: Annotation of islet-associated macrophages based on single-cell analysis.**

**(A)** UMAP embedding of islet-associated macrophages from this study overlaid with cell type annotations from Zakharov et al. [29] following the label transfer workflow.

**(B)** Heatmap showing scaled expression levels of marker genes in each macrophage subpopulation from **Figure 4A**.

**(C)** Enriched gene ontology (GO) terms for marker genes in Macs-1 (top), Macs-4 (middle) and Macs-5 (bottom). Significance (-$\log_{10}$ adjusted p-value) of the enrichments is shown.

**(D)** Violin plot showing normalized expression level of *Ifnb1* across macrophage subpopulations and across different feeding and age groups. **FDR < 0.01, ****FDR < 0.0001; FDR was calculated using the Wilcoxon rank-sum test.

**(E-G)** Volcano plots showing differentially expressed genes in Macs-2 under different conditions: (**E**) WD versus chow at week-1, (**F**) WD versus chow at week-12, and (**G**) aging versus adult (combined week-1 and week-12 chow data). Genes with significant upregulation and downregulation are highlighted in red and blue, respectively. Dashed vertical lines indicate an absolute average $\log_2$-fold change threshold of 0.25, while the dashed horizontal line represents a significance threshold of –log10(adjusted p-value) = 1.3 (corresponding to adjusted p-value of 0.05).

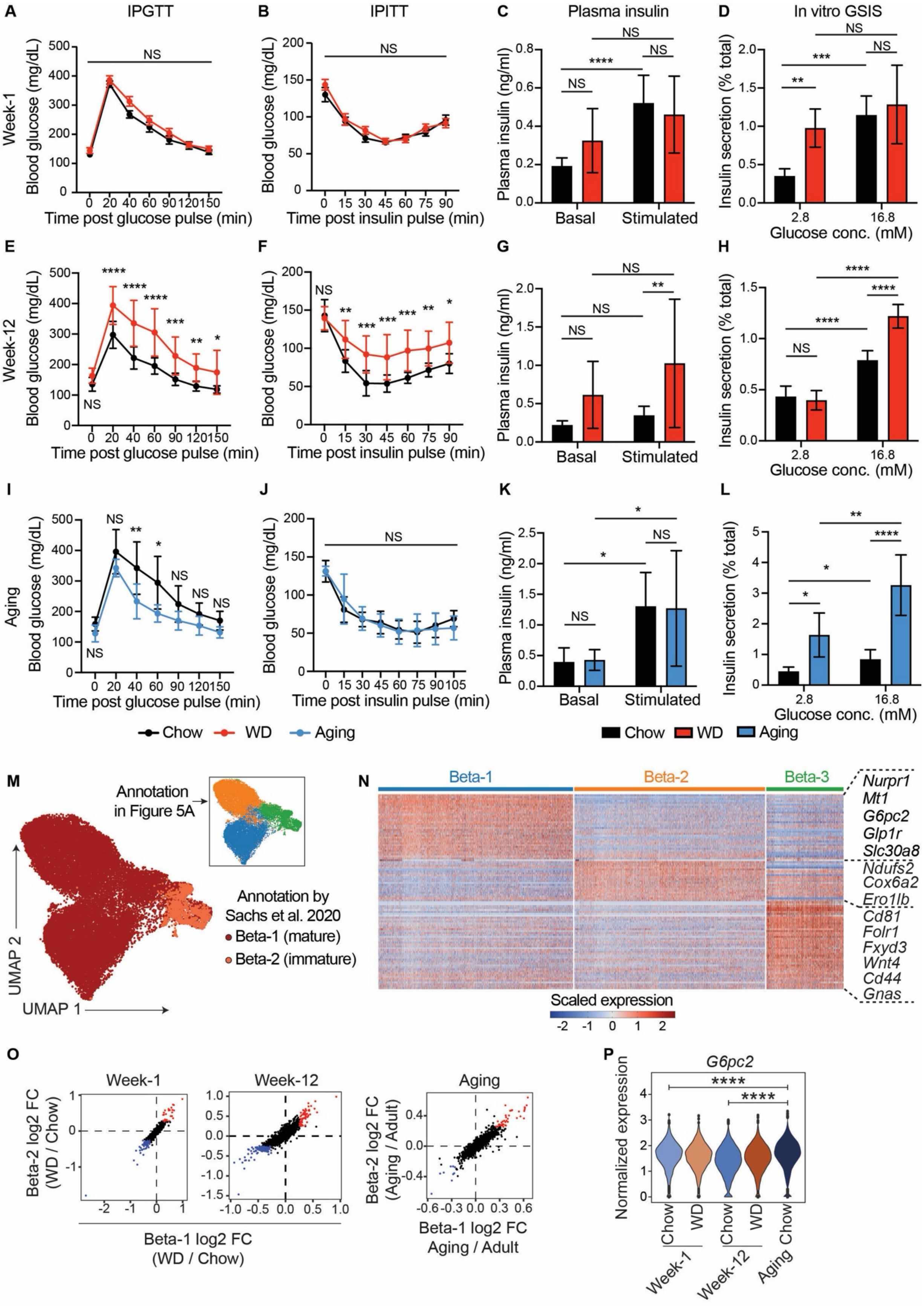
A
IPGTT
B
IPITT
C
Plasma insulin
D
In vitro GSIS
Week-1
Blood glucose (mg/dL)
Time post glucose pulse (min)
Time post insulin pulse (min)
Plasma insulin (ng/ml)
Basal
Stimulated
Insulin secretion (% total)
Glucose conc. (mM)
E
F
G
H
Week-12
I
J
K
L
Aging
Chow
WD
Aging
M
Annotation in Figure 5A
Annotation by Sachs et al. 2020
Beta-1 (mature)
Beta-2 (immature)
UMAP 2
UMAP 1
N
Beta-1
Beta-2
Beta-3
Nurpr1
Mt1
G6pc2
Glp1r
Slc30a8
Ndufs2
Cox6a2
Ero1lb
Cd81
Folr1
Fxyd3
Wnt4
Cd44
Gnas
Scaled expression
O
Week-1
Week-12
Aging
Beta-2 log2 FC (WD / Chow)
Beta-1 log2 FC (WD / Chow)
Beta-2 log2 FC (Aging / Adult)
Beta-1 log2 FC Aging / Adult
P
G6pc2
Normalized expression
Week-1
Week-12
Aging

**Supplementary Figure 5: Functional and transcriptional changes in beta cells in response to western diet and aging.**

**(A, E, I)** Glucose tolerance tests in mice fed chow or WD for 1 week (**A,** n=10 both groups), 12 weeks (**E,** n=10 both groups) or with aging (**I,** adult <3-month-old versus 2-year-old, n=10 both groups). NS, not significant, *p < 0.05, **p < 0.01, ***p < 0.001, ****p < 0.0001 by multiple comparisons test post two-way ANOVA.

**(B, F, J)** Insulin tolerance tests in mice fed chow or WD for 1 week (**B,** n=10 both groups), 12 weeks (**F,** n=10 both groups) or with aging (**J,** adult <3-month-old versus 2-year-old, n=10 both groups). NS, not significant, *p < 0.05, **p < 0.01, ***p < 0.001 by multiple comparisons test post two-way ANOVA.

(**C, G, K**) Plasma insulin levels in mice pre- and post-glucose injection in mice fed chow or WD for 1 week (**C,** n=10 both groups), 12 weeks (**G,** n=10 both groups) or with aging (**K,** adult <3-month-old versus 2-year-old, n=10 both groups). NS, not significant, *p < 0.05, **p < 0.01, ****p < 0.0001 by multiple comparisons test post two-way ANOVA.

**(D, H, L)** Glucose stimulated insulin secretion (GSIS) assay in mouse islets from mice fed chow or WD for 1 week (**D,** 6 mice, with 10 islets collected per mouse for each data point), 12 weeks (**H,** n=6 with 10 islets per data point) or with aging (**L,** adult <3-month-old versus 2-year-old, n=6 with 10 islets per data point). NS, not significant, *p < 0.05, **p < 0.01, ****p < 0.0001 by multiple comparisons test post two-way ANOVA.

**(M)** UMAP embedding of beta cells in this study depicting the cell type annotations from Sachs et al. [39] following the label transfer workflow.

**(N)** Heatmap depicting scaled expression levels of marker genes in each beta cell subtypes from **Figure 5A**.

**(O)** Scatter plots showing average $\log_2$-fold changes in gene expression for Beta-1 and Beta-2 cells after 1-week and 12-week WD feeding compared to chow-fed mice, and in aged versus adult mice.

**(P)** Violin plot showing normalized expression level of *G6pc2* in beta cells across western diet feeding conditions and age. ****FDR $< 0.0001$; FDR was calculated using the Wilcoxon rank-sum test.

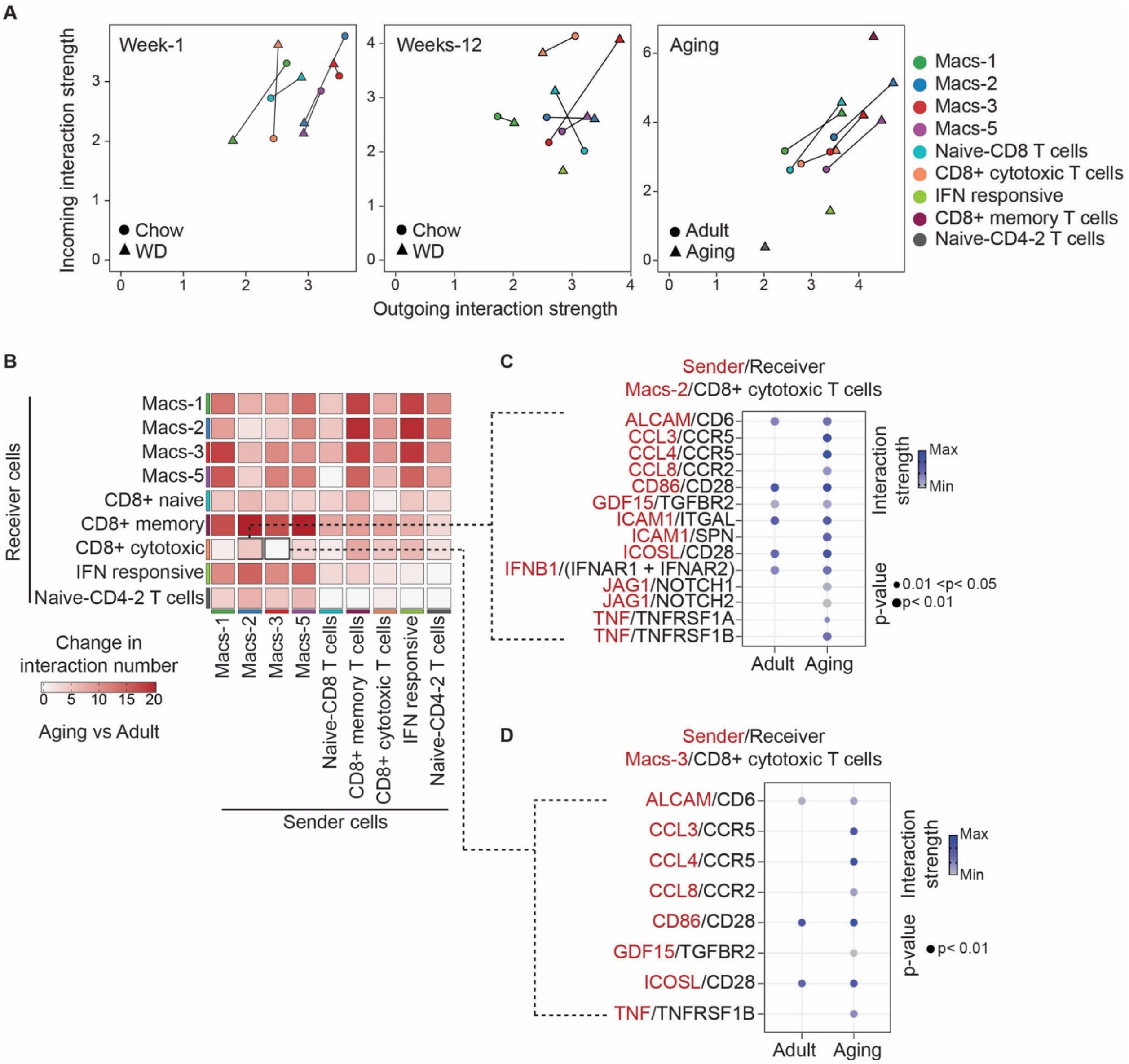


**Supplementary Figure 6: Ligand-receptor analysis in islet-resident immune cells.**

**(A)** Scatter plot showing the outgoing (X-axis) and incoming (Y-axis) interaction strength for the indicated cell populations across various conditions: week-1 chow (n=3 cohorts), week-1 WD (n=2 cohorts), week-12 chow (n=3 cohorts), week-12 WD (n=2 cohorts), aging (n=2 cohorts), and adult (week-1 and week-12 chow, n=6 cohorts).

**(B)** Heatmap showing changes in the number of ligand-receptor pairs among indicated cell populations, comparing adult to aging (n=6 cohorts for adult, n=2 cohorts for aging).

**(C and D)** Dot plots showing the strength of ligand-receptor interactions from Macs-2 to $CD8^+$ cytotoxic T cells **(C)** and from Macs-3 to $CD8^+$ cytotoxic T cells **(D)** in adult and aging conditions. Dot color and size represent the communication probability and p-values, calculated using a one-sided permutation test in CellChat.
